# Selective regulation of kinesin-5 function by β-tubulin carboxy-terminal tails

**DOI:** 10.1101/2024.05.10.593591

**Authors:** Ezekiel C. Thomas, Jeffrey K. Moore

## Abstract

The tubulin code hypothesis predicts that carboxy-terminal tails of tubulins create programs for selective regulation of microtubule-binding proteins, including kinesin motors. However, the molecular mechanisms that determine selective regulation and their relevance in cells are poorly understood. In this work, we report selective regulation of budding yeast kinesin-5 motors, Cin8 and Kip1, by the β-tubulin tail. Cin8, but not Kip1, requires the β-tubulin tail for its recruitment to the mitotic spindle. This creates a balance of both kinesin-5 motors in the spindle, and efficient mitotic progression. We identify a negatively charged patch of acidic amino acids in the β-tubulin tail that mediates the interaction with Cin8 *in vivo*. Using *in vitro* reconstitution with genetically modified yeast tubulin, we demonstrate that the charged patch in the β-tubulin tail increases Cin8 plus-end directed velocity and processivity. Finally, we determine that the positively charged amino-terminal extension from the Cin8 motor domain coordinates interactions with the β-tubulin tail. This work provides the first demonstration of a molecular mechanism underlying selective regulation of closely related kinesin motors by tubulin tails, and how this regulation promotes proper function of the mitotic spindle.

## Introduction

The microtubule cytoskeleton performs various cellular functions, from trafficking to migration to cell division, yet we lack an understanding of how cells generate and interpret molecular differences in microtubule networks to accomplish these functions. The tubulin code hypothesis proposes two means to generate these molecular differences: multiple tubulin isotypes and tubulin post-translational modifications (PTMs) (Janke & Magiera, 2020; Roll-Mecak, 2020). These two facets of the tubulin code intersect at the carboxy-terminal tails (CTTs) of tubulin, which are intrinsically disordered regions that are highly divergent between tubulin isotypes and are hotspots for PTMs. One generally conserved feature of CTTs is their negative charge as a result of enrichment for glutamate and/or aspartate residues. In many eukaryotes, the negative charge of these residues can be further increased by polyglutamylation, a modification where variable chains of glutamate residues are post-translationally ligated to the genetically-encoded glutamates in the CTTs. Polyglutamylation is enriched in specific microtubule structures in axons, mitotic spindles, and axonemes, and thought to create functionally distinct subpopulations of microtubules (Bodakuntla et al., 2020; Lacroix et al., 2010; Redeker et al., 1992; Suryavanshi et al., 2010).

A corollary of the tubulin code hypothesis is that microtubule-associated proteins and motors have divergent sensitivities to tubulin CTTs. Consistent with this notion, microtubule motors do not respond uniformly to changes in CTTs: motors exhibit differential effects on velocity and processivity when measured on microtubules with engineered isotype CTT sequences and modification state (Sirajuddin et al., 2014). Furthermore, kinesin-3 landing rate and motility differs depending on the source of the mammalian tubulin, potentially as a result of differences in isotype composition or polyglutamylation state, and kinesin-1 processivity is affected by polyglutamylation state (Genova et al., 2023; Lessard et al., 2019). However, determining what motors are sensitive to the CTTs and the molecular basis of these interactions remains a major question.

The mitotic spindle presents a compelling case to study these interactions due to division of labor between the subsets of kinetochore, interpolar, and astral microtubules (Winey et al., 1995; Winey & Bloom, 2012). Among these, the interpolar microtubules elongate the mitotic spindle during anaphase to spatially resolve the genome and ensure accurate chromosome inheritance (Cimini et al., 2004; Hoyt et al., 1992; Severin et al., 2001). The budding yeast *Saccharomyces cerevisiae*, like many eukaryotic organisms, accomplishes spindle elongation by generating forces from within the spindle midzone, which is the region of interdigitating, antiparallel microtubules that extend from opposite spindle poles. How these cells generate timely and efficient force generation at a specific region on a subpopulation of microtubules is a fundamental question. Budding yeast have no known modifications to the tubulin CTTs, and a relatively simple repertoire of microtubule-associated proteins and motors that generate force within the midzone. The primary anaphase force generating components of the budding yeast spindle are Kip3 (kinesin-8), Kar3 (kinesin-14), Cin8 and Kip1 (kinesin-5), and Ase1 (MAP65/PRC1). The primary outward force contributing to spindle elongation is generated by the tetrameric kinesin-5 motors that can simultaneously walk on two microtubules, and loss of both Cin8 and Kip1 results in inviable cells (Hoyt et al., 1992; Kapitein et al., 2005; Roof et al., 1992; Saunders & Hoyt, 1992). Although Cin8 and Kip1 have partially redundant roles in spindle elongation, careful examination of null mutants in either gene suggests unique roles: Cin8 plays a more important role in bi-polar spindle assembly and is necessary for the initial, fast phase of anaphase spindle elongation while Kip1 supports the later, slower phase of spindle elongation and stabilizes the late anaphase spindle (Fridman et al., 2013; Leary et al., 2019; Straight et al., 1998). Both Cin8 and Kip1 are also capable of bidirectional motility, adding an extra dimension to the regulation of their motility (Fridman et al., 2013; Gerson-Gurwitz et al., 2011; Roostalu et al., 2011). As such, the budding yeast mitotic spindle provides a system for delineating how different motors use a single CTT state to generate complex function.

Our past work identified distinct roles for the α- or β-CTTs in the budding yeast microtubule network, with the β-CTT specifically promoting spindle assembly and anaphase spindle elongation (Aiken et al., 2014). In addition, we identified a requirement for β-CTT for the localization of Cin8 to the spindle (Aiken et al., 2014). In this study, we first ask which force-generating motors and MAPs in the mitotic spindle are sensitive to the β-CTT. Using *in vivo* fluorescence microscopy, we show that the two kinesin-5 motors Cin8 and Kip1 have differential responses to the loss of the β-CTT. We assign this response to a patch of acidic amino acids within the β-CTT that directly promotes Cin8 function. Using *in vivo* timelapse microscopy and a reconstituted *in vitro* system with Cin8 on yeast microtubules with genetically edited CTTs, we demonstrate the β-CTT promotes the plus-end motility of Cin8. Finally, we determine this interaction is mediated by a basic amino-terminal region extending from the Cin8 motor domain, a feature that may represent an important point of regulation across kinesin motors. These results have implications for kinesin-5 function in more complex systems with multiple β-tubulin isotypes and post-translational modifications such as polyglutamylation, and broadly advance our understanding of how the tubulin-CTTs selectively regulate kinesin function.

## Results

### The β-CTT Differentially Regulates the Budding Yeast Kinesin-5 Motors

Based on our past work that identified the β-CTT, but not the α-CTT, as important for spindle elongation in budding yeast (Aiken et al., 2014), we hypothesized that the β-CTT must regulate force-generating spindle motors and MAPs. We genetically deleted the β-CTT by removing the last 27 codons from chromosomal β-tubulin/*TUB2*, creating *tub2-Δ430*, and predicted that motors and MAPs sensitive to the loss of the β-CTT would have spindle localization levels in these cells that differ from that of wild-type cells. We fluorescently tagged each of the force-generating motors and MAPs in the budding yeast spindle (Kar3, Ase1, Kip3, Cin8, and Kip1) at the endogenous loci and individually quantified the levels of each protein in the pre-anaphase spindle, compared to wild-type *TUB2* cells (Figure 1A). Our previous cryo-electron tomography results indicated an increase in spindle microtubules in *tub2-Δ430* cells, although most of these were very short microtubules (Fees et al., 2016). As a control for microtubule number we measured the amount of the plus-end binding Bik1/CLIP170, which did not display a significant difference in localization to the spindle (*TUB2* = 1.01 ± 0.07, *tub2-Δ430* = 1.10 ± 0.16, p = 0.23; Supplemental Figure 1). Kar3/kinesin-14 (mean ± 95% confidence interval, *TUB2* = 1.03 ± 0.08, *tub2-Δ430* = 1.07 ± 0.24, p = 0.67) and Ase1/MAP65 (*TUB2* = 1.04 ± 0.12, *tub2-Δ430* = 0.95 ± 0.30, p = 0.48) exhibit similar levels in wild-type and *tub2-430*Δ pre-anaphase spindles, suggesting that Kar3 and Ase1 are not sensitive to the loss of the β-CTT (Figure 1B,C). The kinesin-8 Kip3 does display slight decrease in spindle localization in the absence of the β-CTT (*TUB2* = 1.01 ± 0.10, *tub2-Δ430* = 0.80 ± 0.14, p = 0.01; Figure 1D), although we also observed an increase of Kip3 on astral microtubules in the cytoplasm (Supplemental Figure 1B). These results suggest that β-CTT does not promote the localization of Kar3 and Ase1 to the spindle but may enhance the localization of Kip3.

**Figure 1:**
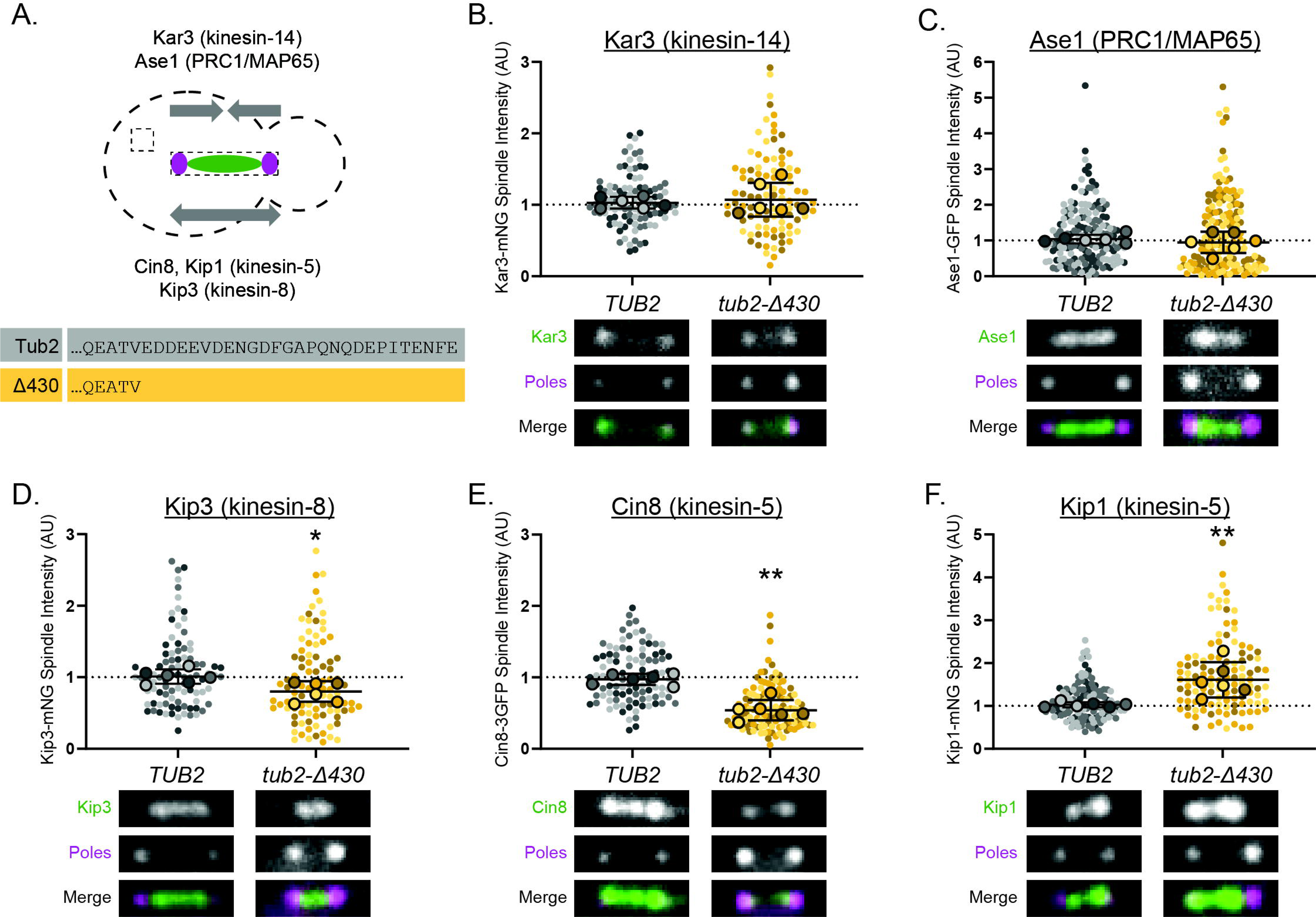
The β-CTT differentially regulates budding yeast kinesin-5 motors. **(A)** Top: Cartoon model depicting the force contribution of spindle midzone components and methods used to quantify spindle localization. Kar3 and Ase1 are thought to contribute inward forces, while Kip3, Cin8, and Kip1 outward forces. A box was draw around the spindle as defined by spindle pole body fluorescence (magenta) and the spindle fluorescence intensity on the spindle was quantified (green). A second box in the cytoplasm was used for background subtraction. Bottom: The amino acid sequence of the Tub2 CTT, which starts at E431, and the deletion in the *tub2-Δ430* allele. **(B)** Quantification (top) and example image (bot) of Kar3-mNeonGreen background-subtracted spindle fluorescence in the presence or absence of the β-CTT. *TUB2* n = 102 cells, *tub2-Δ430* n = 99 cells, p = 0.67. **(C)** Quantification (top) and example image (bot) of Ase1-GFP background-subtracted spindle fluorescence in the presence or absence of the β-CTT. *TUB2* n = 172 cells, *tub2-Δ430* n = 162 cells, p = 0.48. **(D)** Quantification (top) and example image (bot) of Kip3-mNeonGreen background-subtracted spindle fluorescence in the presence or absence of the β-CTT. *TUB2* n = 90 cells, *tub2-Δ430* n = 96 cells, p = 0.01. **(E)** Quantification (top) and example image (bot) of Cin8-3GFP background-subtracted spindle fluorescence in the presence or absence of the β-CTT. *TUB2* n = 104 cells, *tub2-Δ430* n = 117 cells, p < 0.0001. **(F)** Quantification (top) and example image (bot) of Kip1-mNeonGreen background-subtracted spindle fluorescence in the presence or absence of the β-CTT. *TUB2* n = 136 cells, *tub2-Δ430* n = 116 cells, p = 0.0049. The spindle poles in all cells are labeled with Spc110-tdTomato. For each graph in B-F, values are normalized to the median wild-type value. Bolded, outlined points represent the median for each replicate. Error bars are the mean ± 95% CI for the replicate medians. Statistics are student’s t-test between the replicate medians for each protein compared to wild-type. Width of example images = 2.5 µm.

We observed the greatest effects of β-CTT on the kinesin-5 motors Cin8 and Kip1 (Figure 1E, F). Cin8 spindle localization decreased by approximately 50% in the absence of β-CTT (*TUB2* = 0.97 ± 0.08, *tub2-Δ430* = 0.54 ± 0.14; p < 0.0001), while Kip1 increased by approximately 50% (*TUB2* = 1.03 ±0.06, *tub2-Δ430* = 1.61 ± 0.41; p = 0.0049). While the decrease in Cin8 spindle localization is consistent with our past results (Aiken et al., 2014), the increase in Kip1 localization was unexpected, since the two motors are from the same kinesin-5 family. Overall, these results suggest that the β-CTT predominantly regulates the kinesin-5 motors Cin8 and Kip1 to promote spindle elongation.

### Comparing the Expression and Spindle Localization of Kinesin-5 Motors

Because β-CTT appears to have opposite effects on Cin8 and Kip1, we next wanted to determine whether this could be attributed to differences in expression during the cell cycle or localization to distinct sub-regions of the spindle. Previous studies have established that Cin8 plays an important role during spindle assembly and the fast phase of anaphase spindle elongation, while Kip1 plays an important role later in anaphase to stabilize the spindle (Fridman et al., 2013; Leary et al., 2019; Straight et al., 1998). These data suggest that Cin8 may localize to the spindle earlier than Kip1, so we predicted that Cin8 may be expressed earlier in the cell cycle. To test this prediction, we added carboxy-terminal 3HA tags to native Cin8 and Kip1, arrested cells in G1, then synchronously released cells and collected samples every 15 minutes for western blot analysis. Cin8 and Kip1 exhibit similar patterns of expression after G1 release – increasing at approximately 30 minutes, peaking between 45 and 60 minutes, and decreasing between 75 and 90 minutes (Figure 2A,B). This represents increase during S-phase, peak during G2/M, and decrease during mitotic exit. Although the cell-cycle timing of Cin8 and Kip1 expression is similar, our results show that cells contain approximately 2X more Cin8 than Kip1 during mitosis (Figure 2A,B).

**Figure 2:**
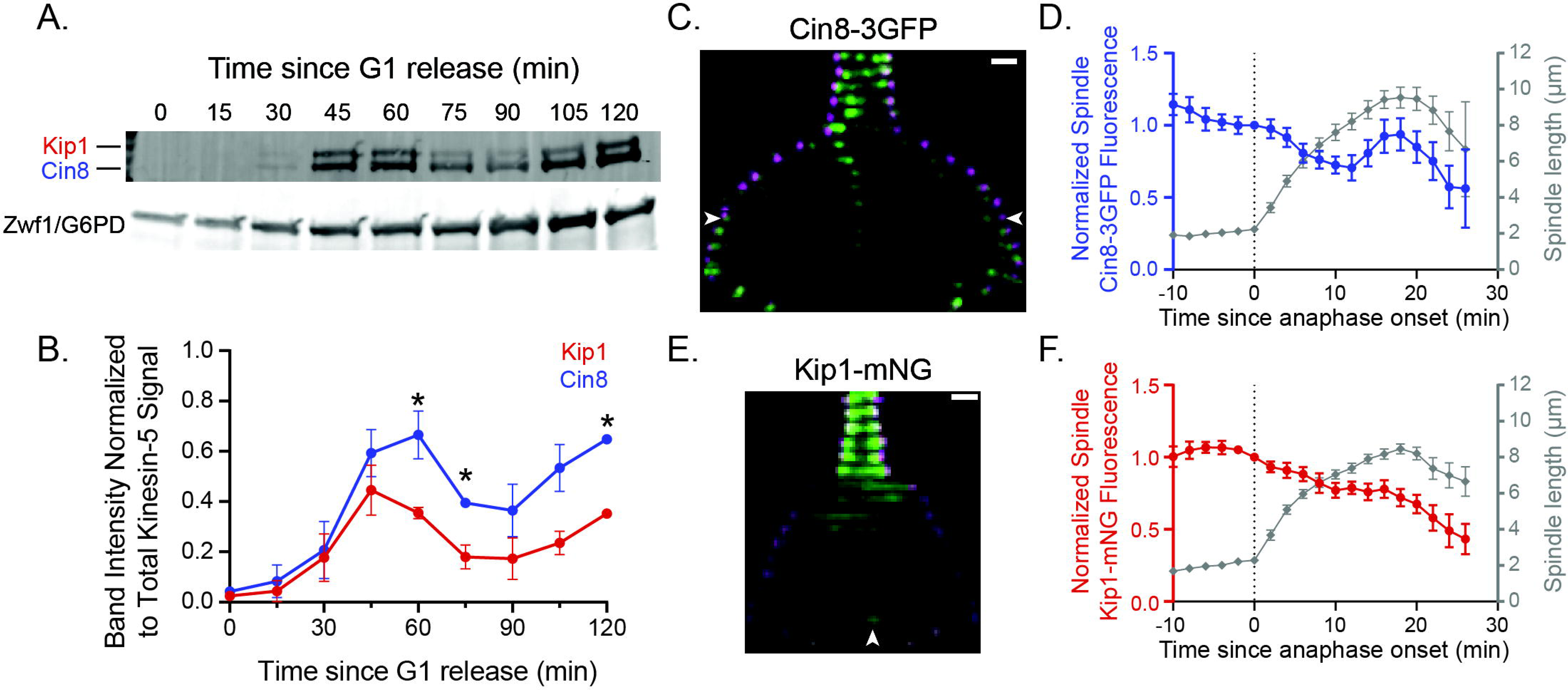
Comparing kinesin-5 expressing and spindle localization. **(A)** Example western blot against Cin8-3HA (121 kDa) and Kip1-3HA (130 kDa) or Zwf1/Glucose-6-phosphate dehydrogenase loading control. Cells were arrested in G1, released, and lysate collected every 15 minutes. **(B)** Quantification of time course western blots against Cin8-3HA and Kip1-3HA such as in panel A. Anti-HA intensity normalized to Zwf1 loading control and then normalized to total anti-HA signal at 120 minutes. Error bars are standard deviation. n = 2 independent experiments; * p < 0.05 by student’s t-test. **(C)** Example montage of cells expressing Cin8-3GFP (green) and Spc110-tdTomato (magenta). Arrowheads point out Cin8 localization adjacent to the spindle poles during late anaphase. First timepoint is 10 minutes prior to anaphase onset. Time interval = 2 min. Scale bar = 1 µm. **(D)** Quantification of background-subtracted Cin8-3GFP spindle fluorescence intensity (left axis) and spindle length (right axis) as a function of time since anaphase onset. Cin8-3GFP intensity normalized to anaphase onset. Error bars are mean ± 95% CI. n = 21 cells. **(E)** Example montage of cells expressing Kip1-mNeonGreen (green) and Spc110-tdTomato (magenta). Arrowhead points out Kip1 localization at the middle of the spindle during late anaphase. First timepoint is 10 minutes prior to anaphase onset. Time interval = 2 min. Scale bar = 1 µm. **(F)** Quantification of background-subtracted Kip1-mNeonGreen spindle fluorescence intensity (left axis) and spindle length (right axis) as a function of time since anaphase onset. Kip1-mNeonGreen intensity normalized to anaphase onset. Error bars are mean ± 95% CI. n = 24 cells.

To quantify changes in kinesin-5 localization during the cell cycle, we released G1 arrested cells and every 2 minutes imaged spindle poles labeled with Spc110-tdTomato along with either Cin8-3GFP or Kip1-mNeonGreen, all tagged at the native loci (Figure 2C-F). We quantified kinesin-5 signal in the spindle and normalized to the level at anaphase onset, as determined by change in spindle length over time. Spindle localization for both Cin8 and Kip1 peaks prior to anaphase onset and then decreases as cells enter anaphase (Figure 2C-F). This decrease at anaphase onset was not due to photobleaching; we found examples of cells entering anaphase up to 16 minutes apart with similar levels of Cin8 or Kip1 signal on the spindle prior to anaphase onset. There are, however, differences the spindle localization of either kinesin-5 during late anaphase. Cin8 levels decrease during early anaphase and then increase during late anaphase (Figure 2C,D). This increase in Cin8 levels during late anaphase represents a re-localization of Cin8 to the spindle poles prior to spindle disassembly (see arrowheads in Figure 2C). In contrast, Kip1 levels decrease during early anaphase, and then remain consistent during late anaphase (Figure 2E,F). During late anaphase, we noticed that Kip1 is enriched in the middle of the spindle (arrowhead Figure 2E). This is in agreement with the past results suggesting a role in stabilizing the late-anaphase spindle (Fridman et al., 2013). Overall, these results indicate that the timing of Cin8 and Kip1 expression and localization to the spindle is similar, but Cin8 is more abundant than Kip1 during mitosis and re-localizes to a different region of the spindle in late anaphase.

### Mitotic Delay Increases Kip1 Levels

We hypothesized three non-mutually exclusive models to explain the decrease in Cin8 and increase in Kip1 spindle localization in the absence of the β-CTT (Figure 3A). 1) Cin8 and Kip1 compete for access to binding sites on the mitotic spindle, which could cause an inverse relationship in their spindle localization. 2) A direct interaction between the β-CTT and the kinesin-5 motors with the β-CTT promoting Cin8 localization and/or inhibiting Kip1 localization. 3) The β-CTT promotes mitotic progression, which has an indirect effect on Cin8 and Kip1 levels. To test the first model, we predicted that knocking out one kinesin-5 would allow increased localization of the other to the spindle. To test this possibility, we imaged asynchronous *kip1Δ* cells expressing Cin8-3GFP or *cin8Δ* cells expressing Kip1-mNeonGreen. Cin8 spindle localization slightly increases in *kip1Δ* cells compared to wild type (mean ± 95% confidence interval, *KIP1* = 1.01 ± 0.15, *kip1Δ* = 1.18 ± 0.10, p = 0.03; Figure 3B), while Kip1 levels shows a stronger increase in *cin8Δ* cells compared to wild type (*CIN8* = 0.97 ± 0.15, *cin8Δ* = 1.57 ± 0.35, p = 0.002; Figure 3C). The level of increase for Kip1 in *cin8Δ* cells was reminiscent of the increased Kip1 localization in the absence of the β-CTT (Figure 1F). These results suggest that Cin8 and Kip1 may compete for limited binding sites within the spindle.

**Figure 3:**
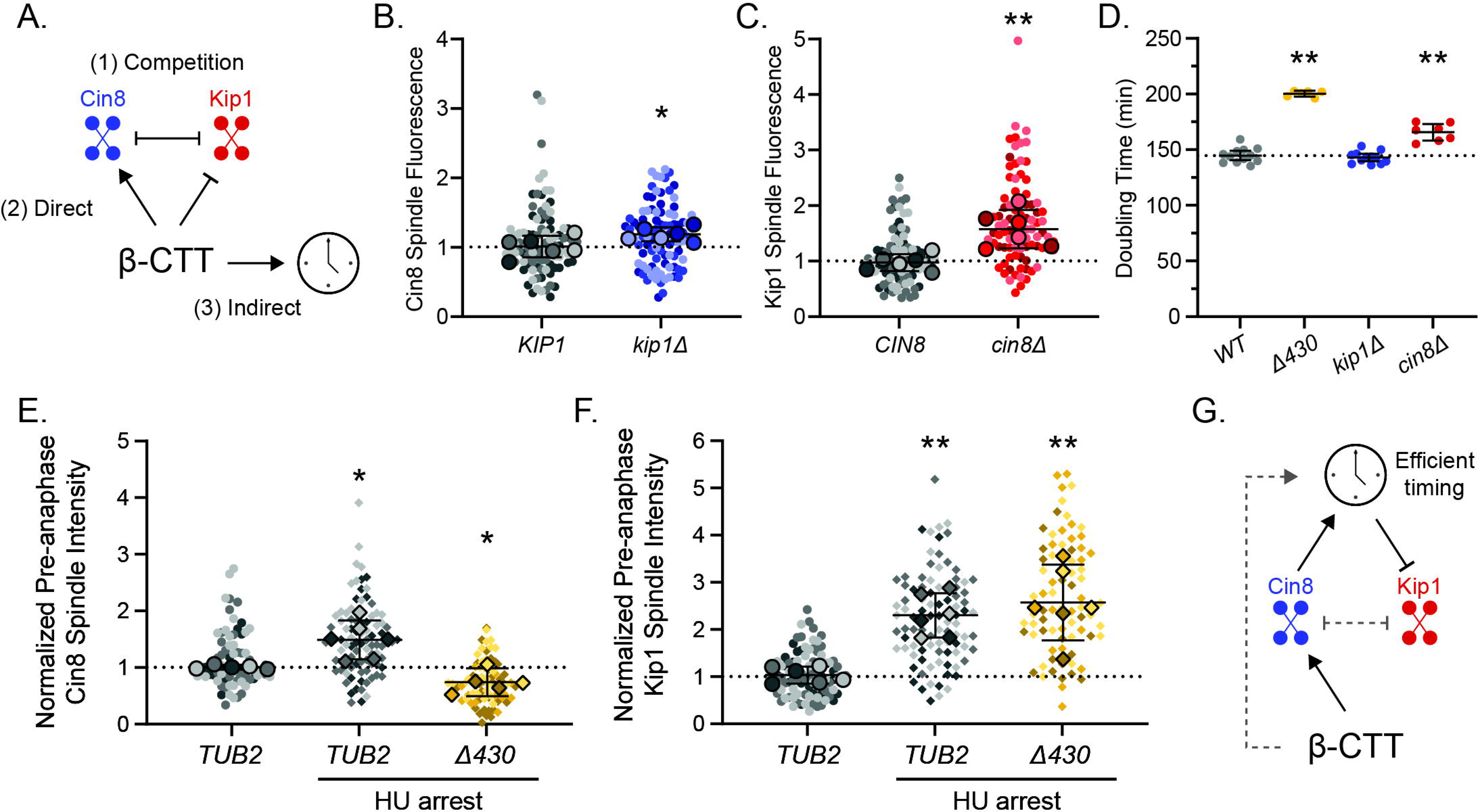
Mitotic delay increases Kip1 levels. **(A)** Cartoon depicting possible non-mutually exclusive models to explain the opposite response of Cin8 and Kip1 to the loss of the β-CTT. **(B)** Quantification of background-subtracted Cin8-mNeonGreen spindle fluorescence in wild-type *KIP1* or *kip1Δ* cells. Values are normalized to the median wild-type value. Bolded, outlined points represent the median for each replicate. Error bars are the mean ± 95% CI for the replicate medians. *KIP1* n = 95 cells, *kip1Δ* n = 97 cells. p = 0.03; statistics are student’s t-test for the replicate medians. **(C)** Quantification of background-subtracted Kip1-mNeonGreen spindle fluorescence in wild-type *CIN8* or *cin8Δ* cells. Values are normalized to the median wild-type value. Bolded, outlined points represent the median for each replicate. Error bars are the mean ± 95% CI for the replicate medians. *CIN8* n = 93 cells, *cin8Δ* n = 89 cells. p = 0.002; statistics are student’s t-test for the replicate medians. **(D)** Doubling time in minutes for wild-type (WT), *tub2-Δ430* (p < 0.0001, t-test with Welch’s correction), *kip1Δ* (p = 0.46, t-test), or *cin8Δ* (p = 0.0001, t-test with Welch’s correction) cells. Error bars are the mean ± 95% CI. Statistics are compared to wild type. **(E)** Quantification of background-subtracted Cin8-3GFP spindle fluorescence in asynchronous *TUB2* cells or two-hour treatment with hydroxyurea (HU) to arrest *TUB2* or *tub2-Δ430* cells. Values are normalized to the median asynchronous wild-type value. Bolded, outlined points represent the median for each replicate. Error bars are the mean ± 95% CI for the replicate medians. Asynchronous *TUB2* n = 75 cells, HU-treated *TUB2* n = 89 cells, HU-treated *tub2-Δ430* n = 78 cells. Compared to asynchronous *TUB2*, HU-treated *TUB2* p = 0.02, HU-treated *tub2-Δ430* p = 0.04; student’s t-test with Welch’s correction. **(F)** Quantification of background-subtracted Kip1-mNeonGreen spindle fluorescence in asynchronous *TUB2* cells or two-hour treatment with hydroxyurea (HU) to arrest *TUB2* or *tub2-Δ430* cells. Values are normalized to the median asynchronous wild-type value. Bolded, outlined points represent the median for each replicate. Error bars are the mean ± 95% CI for the replicate medians. Asynchronous *TUB2* n = 88 cells, HU-treated *TUB2* n = 93 cells, HU-treated *tub2-Δ430* n = 80 cells. Compared to asynchronous *TUB2*, HU-treated *TUB2* p = 0.0005, HU-treated *tub2-Δ430* p = 0.004; student’s t-test with Welch’s correction. **(G)** Proposed model for how the β-CTT regulates Cin8 and Kip1. The β-CTT directly promotes Cin8 function, which in turn promotes efficient mitotic timing and inhibits excessive Kip1 spindle localization. The grey dotted lines represent our data do not rule out the possibility that the β-CTT promotes efficient mitotic timing through other factors such as microtubule dynamics. Cin8 and Kip1 may also inhibit each other’s spindle localization by competing for binding sites on the spindle.

Our past work demonstrated that mutant cells lacking β-CTT experience mitotic delay and depend on the spindle assembly checkpoint for successful mitosis (Aiken et al., 2014; Fees et al., 2016). Because of these data, we hypothesized that cells lacking one of the two kinesin-5 motors may experience a mitotic delay. First, we measured the doubling time of *tub2-Δ430* or kinesin-5 knock-out cells. In agreement with our past results, cells lacking the β-CTT exhibited a slower doubling time, indicating a growth delay (wild type: 140.5 ± 4.3 min, mean ± 95% confidence interval, *tub2-Δ430*: 200.3 ± 2.8 min, p < 0.0001; Figure 3D). Interestingly, *kip1Δ* cells did not display an increase in doubling time (142.9 ± 3.3 min), but *cin8Δ* cells did have a delay (165.7 ± 7.3 min, p = 0.0001 compared to wild type). These results are consistent with the idea that Cin8 promotes efficient mitotic progression (Hoyt et al., 1992; Saunders & Hoyt, 1992; Straight et al., 1998) and suggest that a cell cycle delay in *cin8Δ* cells may also contribute to an increase in Kip1 spindle localization. To further test the mitotic timing model, we next asked if delaying mitotic progression would affect the spindle localization of Cin8 and Kip1. We predicted an increase in Kip1 spindle localization in arrested cells because Kip1 spindle localization increased in *tub2-Δ430* cells and *cin8Δ* cells, which both exhibit an increased doubling time. We treated asynchronous log-phase cells with hydroxyurea for 2 hours to arrest the cells in S phase with short bipolar spindles and quantified the spindle localization of either Cin8 or Kip1. In arrested wild-type cells, spindle localization of Cin8 and Kip1 increased relative to asynchronous wild-type cells (Cin8 levels: asynchronous mean = 1.01 ± 0.04, arrested = 1.49 ± 0.34, p = 0.02; Kip1 levels: asynchronous = 1.03 ± 0.18, arrested = 2.30 ± 0.47, p = 0.0005; Figure 3E,F). However, arrested cells that lack the β-CTT still exhibit a reduced amount of Cin8 localizing to the spindle (0.74 ± 0.25, p = 0.04 compared to asynchronous *TUB2* cells; Figure 3E). This result suggests that despite the increased Cin8 spindle localization in arrested wild-type cells, arrested cells still require the β-CTT to promote Cin8 spindle localization. Contrastingly, Kip1 levels in arrested *tub2-Δ430* cells were increased relative to asynchronous wild-type *TUB2* cells but not statistically different from arrested wild-type cells (2.57 ± 0.80, p = 0.004 compared to asynchronous *TUB2* cells, p = 0.48 compared to arrested *tub2-Δ430* cells; Figure 3F). The fact that Kip1 levels are similarly increased in *TUB2* and *tub2-Δ430* cells indicates arresting cells with a bipolar spindle is sufficient to increase Kip1 spindle localization independent of the β-CTT, arguing against the β-CTT directly regulating Kip1. We propose the β-CTT directly promotes Cin8 function, which in turn promotes efficient mitotic progression and attenuates levels of Kip1 on the spindle (Figure 3G).

### The β-CTT Acidic Patch Promotes Cin8 Spindle Localization and Recruits Cin8 to the Midzone

We next sought to identify the features of the β-CTT that regulate Cin8. A common characteristic of the tubulin CTTs is their negative charge, owing to the enrichment of glutamate and aspartate residues. The charge from these side chains is thought to mediate interactions with microtubule-associated proteins. The budding yeast β-CTT includes a patch of acidic amino acids (residues 431 to 438; Figure 4A) that represents more than half of the acidic residues within the β-CTT. We predicted that if the negative charge of the β-CTT is sufficient to promote Cin8 function, returning the acidic patch (*tub2-Δ438*) would restore wild-type Cin8 spindle levels compared to a neutrally charged patch (*tub2-Δ438Q*; Figure 4A). To test this prediction, we quantified the pre-anaphase spindle localization of Cin8 in asynchronous cells expressing these β-CTT alleles (Figure 4B). The β-CTT acidic patch is sufficient to support wild-type levels of Cin8 localization to the spindle (*TUB2* = 1.03 ± 0.08, *tub2-Δ438* = 1.00 ± 0.14, p = 0.66). In contrast, the neutral patch allele *tub2-Δ438Q* exhibits significantly decreased Cin8 localization, but at a level that is greater than in *tub2-Δ430* cells (*tub2-Δ438Q* = 0.86 ± 0.09, p = 0.005 compared to *TUB2*; p < 0.001 compared to *tub2-Δ430;* Figure 4B).

**Figure 4:**
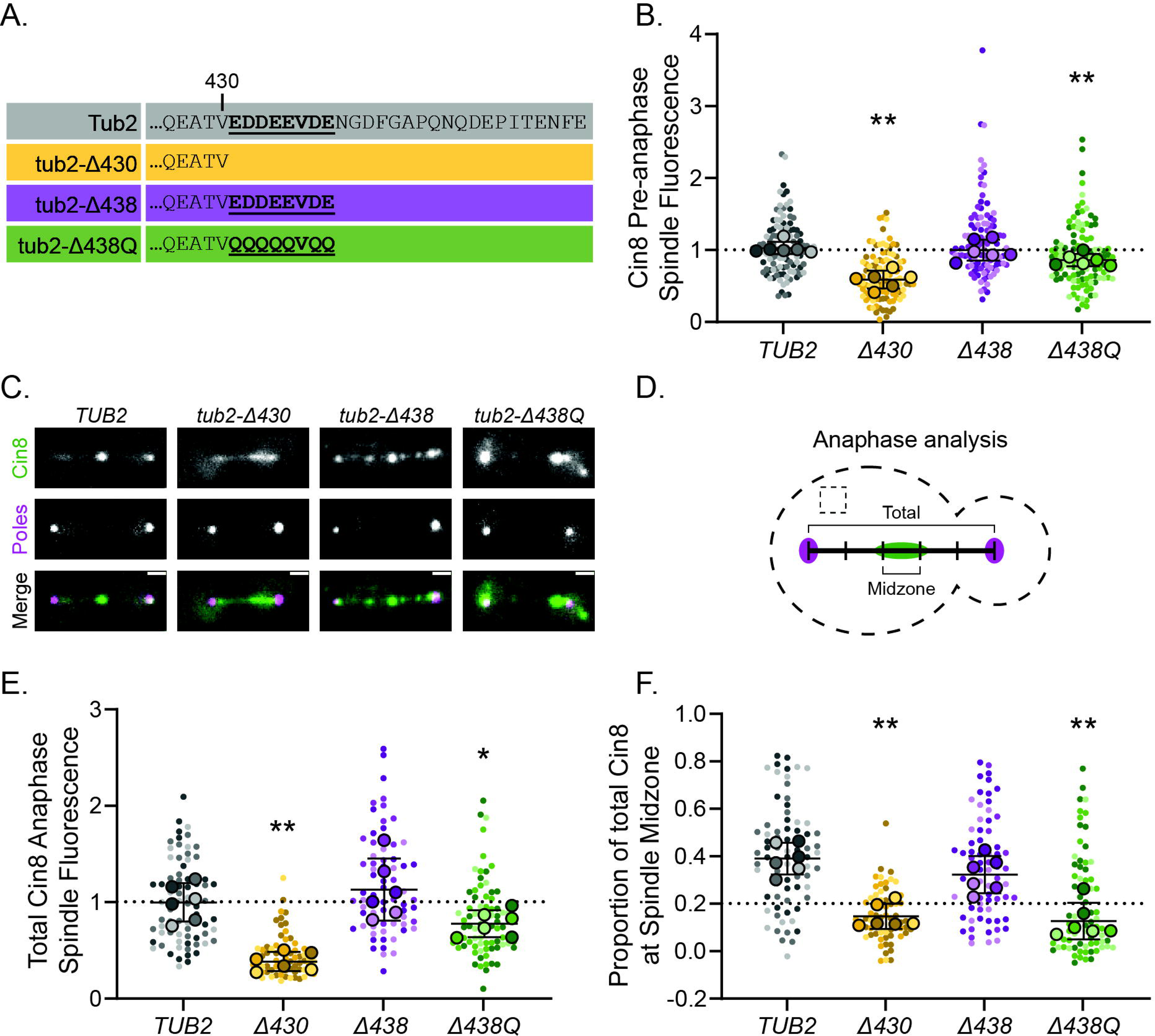
The β-CTT acidic patch promotes Cin8 spindle localization and recruits Cin8 to the midzone. **(A)** Table of wild-type Tub2 and mutant tub2 β-CTT amino acids. The bold, underlined residues represent the “acidic patch,” E431 through E438. **(B)** Quantification of background-subtracted Cin8-3GFP pre-anaphase spindle fluorescence in wild-type *TUB2* or mutant *tub2* cells. Values are normalized to the median wild-type value. Bolded, outlined points represent the median for each replicate. Error bars are mean ± 95% CI for the replicate medians. *TUB2*: n = 113 cells; *tub2-Δ430*: n = 113 cells, p < 0.0001; *tub2-Δ438*: n = 112 cells, p = 0.66; *tub2-Δ438Q*: n = 117 cells, p = 0.005. Statistics are student’s t-test compared to wild-type *TUB2*. **(C)** Example anaphase spindle images of Cin8-3GFP and Spc110-tdTomato (poles) in wild-type *TUB2* or mutant *tub2* cells. Scale bars = 1 µm. **(D)** Cartoon model of methods to quantify anaphase spindle localization. A 3-pixel wide line scan was drawn along the spindle as determined by Cin8-3GFP and Spc110-tdTomato fluorescence, and a box in the cytoplasm was used to determine the cellular background for subtraction. The length of the spindle was divided into five equally sized bins and the background-subtracted intensity within each bin along the line scan was calculated. **(E)** Quantification of background-subtracted Cin8-3GFP anaphase spindle fluorescence along the total length of the spindle in wild-type *TUB2* or mutant *tub2* cells. Values are normalized to the median wild-type value. Bolded, outlined points represent the median for each replicate. Error bars are mean ± 95% CI for the replicate medians. *TUB2*: n = 76 cells; *tub2-Δ430*: n = 69 cells, p < 0.0001; *tub2-Δ438*: n = 71 cells, p = 0.39; *tub2-Δ438Q*: n = 75 cells, p = 0.04. Statistics are student’s t-test compared to wild-type *TUB2*. **(F)** Quantification of proportion of background-subtracted Cin8-3GFP anaphase spindle fluorescence in the middle 20% of the spindle in the same cells from G. Values are normalized to the median wild-type value. Bolded, outline pointed represent the median for each replicate. Error bars are mean ± 95% CI for the replicate medians. *TUB2*: n = 76 cells; *tub2-Δ430*: n = 69 cells, p < 0.0001; *tub2-Δ438*: n = 71 cells, p = 0.12; *tub2-Δ438Q*: n = 75 cells, p < 0.0001. Statistics are student’s t-test compared to wild-type *TUB2*.

Kip1 exhibits similar localization in wild-type *and tub2-Δ438* cells (*TUB2* = 1.02 ± 0.09, *tub2-Δ438* = 1.05 ± 0.42, p = 0.87), and similarly increased localization in both the neutral patch mutant in the absence of the β-CTT (*tub2-Δ430* = 1.56 ± 0.38, *tub2-Δ438Q* = 1.76 ± 0.50, p = 0.01 and p = 0.01, respectively compared to *TUB2*; Supplemental Figure 2). These results suggest that the β-CTT acidic patch is sufficient for Cin8 spindle localization, and again support a model in which Kip1 levels are elevated as a secondary consequence of Cin8 disruption.

We next asked whether the β-CTT is important for Cin8 localization to the spindle during anaphase. Cin8 forms bright puncta along anaphase spindles in wild-type cells, compared to weaker and diffuse spindle localization in *tub2-Δ430* cells (Figure 4C). We also observed increased background GFP fluorescence around the spindle poles in *tub2-Δ430* cells, presumably representing unbound signal in the nucleoplasm (Figure 4C). Both *tub2-Δ438* and *tub2-Δ438Q* cells exhibit puncta of Cin8 along the spindle, although *tub2-Δ438Q* cells show reduced Cin8 signal at the middle of the spindle (Figure 4C). To quantify these differences, we measured Cin8-3GFP signal intensity along a 3 pixel-wide (approximately 160 nm) segmented line connecting the spindle pole in the mother to the spindle pole in the bud (Figure 4D). For this analysis, we only included anaphase cells with a spindle length between 2.5 and 6 µm and with at least some co-linear Cin8 signal to suggest an intact anaphase spindle. The total Cin8 anaphase spindle localization (i.e., the sum of values along the linescan) mirrored the trends of pre-anaphase spindles with Cin8 levels reduced in cells lacking the β-CTT acidic patch (*TUB2* = 0.99 ± 0.20; *tub2-Δ430* = 0.38 ± 0.10, p < 0.0001; *tub2-Δ438* = 1.13 ± 0.32, p = 0.39; *tub2-Δ438Q* = 0.77 ± 0.14, p = 0.04; statistics compared to *TUB2*; Figure 4E).

Because Cin8 appears to be depleted from the middle of the spindle in *tub2-Δ438Q* cells, we measured relative enrichment at the spindle midzone. We defined the midzone as the middle 20% of the anaphase spindle, which roughly corresponds to the region where Ase1 localizes (Figure 4D) (Thomas et al., 2020). In wild-type anaphase cells, 38.9 ± 6.6% of total Cin8 is located at the midzone, which is nearly double the amount that would be expected from an even distribution across quintiles in our analysis (Figure 4F, Supplemental Figure 2). In contrast, *tub2-Δ430* anaphase cells exhibit 14.6 ± 5.3% of total Cin8 in the midzone (p < 0.0001 compared to *TUB2*; Figure 4F). Returning the acidic patch with the *tub2-Δ438* allele restored Cin8 recruitment to the midzone (32.1 ± 7.8%, p = 0.12 compared to *TUB2*; Figure 4F), but the neutral patch failed to enrich Cin8 at the midzone (12.7 ± 7.7%, p < 0.0001 compared to *TUB2*; Figure 4F). It is important to note these values represent the fraction of total Cin8 on the spindle per cell, so decreased total spindle levels alone would not explain these reductions in recruitment to the middle of the spindle. Instead, these results suggest that the β-CTT acidic patch promotes Cin8 localization to the spindle and recruitment to the midzone of the anaphase spindle.

### The β-CTT Promotes Cin8 Plus-End Motility

Based on our results from anaphase spindles, we hypothesized that the β-CTT may promote the motility of Cin8 towards microtubule plus ends. A prediction of this hypothesis would be more frequent, faster, or longer plus-end directed Cin8 motility events. To test these predictions, we used time lapse microscopy of anaphase spindles in living cells and imaged labeled Cin8 and spindle poles every 5 seconds. We then generated kymographs by aligning these image series at the geometric center of the spindle, as determined by spindle pole fluorescence. For this analysis, we compared wild-type cells to mutant cells expressing *tub2-Δ438Q*, since *tub2-Δ430* cells display weak Cin8 signal that is not amenable to time lapse imaging. In wild-type cells most of the Cin8 is enriched at the midzone during anaphase spindle elongation. In contrast, cells expressing *tub2-Δ438Q* display less Cin8 at the midzone and more enrichment closer to the spindle poles (Figure 4F, 5A). In both cases, we observed motility events in which Cin8 traveled toward the middle of the spindle (Figure 5A arrowheads). The velocity of these midzone-directed events in wild-type cells is 19.40 ± 2.57 nm/s, which is similar to past reported values (Figure 5B)(Gerson-Gurwitz et al., 2011). In contrast, Cin8 exhibits slower midzone-directed velocity in *tub2-Δ438Q* cells (13.8 ± 2.85 nm/s, p = 0.0046; Figure 5A,B). We did not observe any clear minus-end directed events, but our imaging conditions may be unable to capture the rapid minus-end directed velocity that has been previously described (Gerson-Gurwitz et al., 2011). These results indicate that the β-CTT promotes the movement of Cin8 towards the midzone during anaphase. However, a caveat of this experiment is that the spindle contains parallel and antiparallel microtubules, making the plus or minus-end directionality of movement difficult to determine.

**Figure 5:**
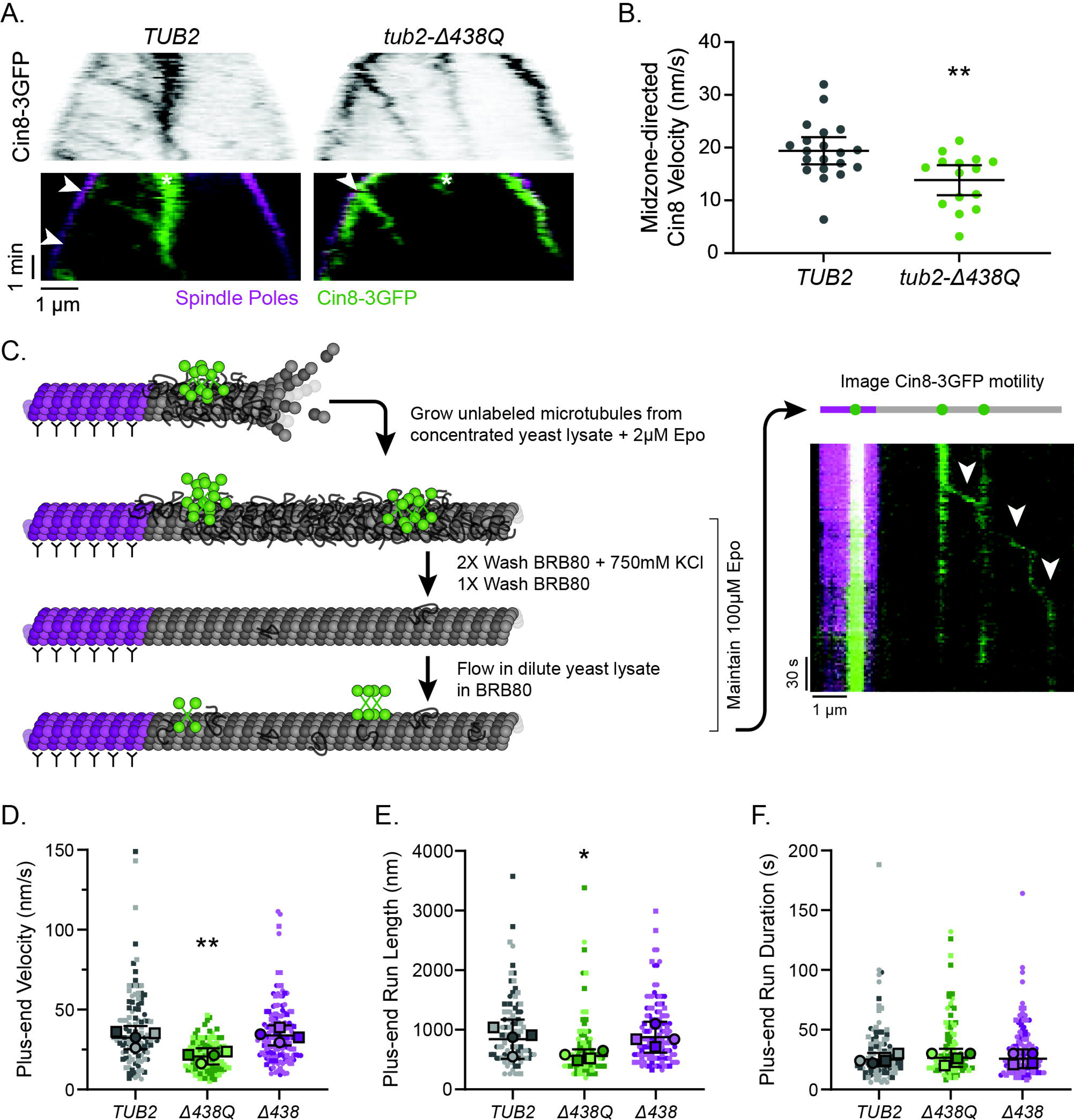
The β-CTT Promotes Cin8 Plus-End Motility. **(A)** Example kymographs from timelapse fluorescence microscopy of Cin8-3GFP and Spc110-tdTomato (poles) in cells at 30°C. Arrowheads represent midzone-directed motility events. Asterisks mark spindle midzone. **(B)** Quantification of velocity of Cin8 midzone-directed motility events in cells expressing *TUB2* (n = 20 events) or *tub2-Δ438Q* (n = 15 events). Error bars are mean ± 95% CI. p = 0.0046; student’s t-test. **(C)** Cartoon model of *in vitro* Cin8-3GFP motility on yeast microtubules. First, a concentrated yeast lysate was used to grow yeast microtubules from rhodamine-labeled, porcine brain GMPCPP seeds. Next, proteins were removed from the grown microtubules with several wash steps. Microtubules were maintained with a high concentration of stabilizing epothilone A (Epo). Finally, a dilute yeast lysate was used to image motility. An example kymograph is shown on the right. **(D)** Quantification of Cin8 plus-end directed velocity. *TUB2*: n = 111 events; *tub2-Δ438Q*: n = 113 events, p = 0.0056; *tub2-Δ438*: n = 149 events, p = 0.67. Statistics are student’s t-test of replicate medians compared to wild-type *TUB2*. Error bars mean ± 95% CI. **(E)** Quantification of Cin8 plus-end distance traveled during motility event from D with a start and stop during the imaging. *TUB2*: n = 89 events; *tub2-Δ438Q*: n = 84 events, p = 0.041; *tub2-Δ438*: n = 119 events, p = 0.81. Statistics are student’s t-test of replicate medians compared to wild-type *TUB2*. Error bars mean ± 95% CI. **(F)** Quantification of time spent moving during a plus-end directed motility event from D with a start and stop during the imaging. *TUB2*: n = 89 events; *tub2-Δ438Q*: n = 84 events, p = 0.63; *tub2-Δ438*: n = 119 events, p = 0.81. Statistics are student’s t-test of replicate medians compared to wild-type *TUB2*. Error bars mean ± 95% CI.

To clearly determine how the β-CTT impacts directional Cin8 movement on microtubules, we shifted to an *in vitro* system. We developed an experiment using yeast cell lysate to examine Cin8 motility on native, homogenous microtubules, instead of tubulin sourced from bovine or porcine brains that is highly heterogenous. Prior studies using yeast cell extracts failed to observe motile Cin8 on native yeast microtubules (Torvi et al., 2022). We speculated this might be because those studies used high concentrations of cell lysates, which could create immotile Cin8 aggregates (Bell et al., 2017; Pandey, Reithmann, et al., 2021). First, using the same yeast strains from our *in vivo* imaging experiments, we made a high concentration lysate mixture (final protein concentration of approximately 1.5 mg/ml) to assemble unlabeled yeast microtubules from rhodamine-labeled GMPCPP porcine brain seeds that were adhered to the coverslip (Figure 5C). The assembled microtubules were stabilized with 2 μM epothilone A. Second, we washed with high ionic strength buffer (750 mM KCl) to remove bound MAPs from the microtubules. Third, we flowed in a lower concentration mixture of the same yeast lysate (final protein concentration of approximately 50 µg/ml) to return labeled Cin8 to the pre-assembled microtubules. With this method we were able to observe processive bidirectional motility of Cin8 foci, as previously reported for Cin8 on brain tubulin (Gerson-Gurwitz et al., 2011; Roostalu et al., 2011). Microtubules were maintained throughout this process as determined by interference reflection microscopy and separate experiments using lysate from GFP-Tub1 expressing cells (Supplemental Figure 3).

Using this system, we tested the prediction that disrupting the charge of the β-CTT would reduce Cin8 plus-end motility. Cin8 plus-end-directed velocity on microtubules polymerized from wild-type *TUB2* cell lysate averaged 32.5 ± 7.3 nm/s (mean ± 95% confidence interval, Figure 5D). Cin8 velocity is similar on microtubules from *tub2-Δ438* cell lysate (33.8 ± 6.3 nm/s, p = 0.67), but slower on microtubules from *tub2-Δ438Q* cell lysate (20.7 ± 5.0 nm/s, p = 0.0056; Figure 5D). We also quantified the run length and duration for all plus-end-directed motility events for which we observed the beginning and end of motility. This analysis shows a significant reduction in the run length of Cin8 on *tub2-Δ438Q* microtubules (*TUB2* = 845 ± 330 nm; *tub2-Δ438Q* = 560 ± 114 nm, p = 0.0411; *tub2-Δ438* = 877 ± 260 nm, p = 0.81 compared to wild type; Figure 5E), but not in the run duration (*TUB2* = 25 ± 5.5 s, *tub2-Δ438Q* = 26.5 ± 7.5 s, *tub2-Δ438* = 25.8 ± 7.9 s; Figure 5F). We also assessed minus-end-directed motility events in our imaging data. The velocities of these events were more variable than plus-end directed motility events, making them difficult to compare across genotypes. Our data suggest that Cin8 minus-end-directed velocity may be reduced on microtubules grown from *tub2-Δ438Q* cell lysate, but these differences are not significant (*TUB2* = 59.1 ± 29.1 nm/s; *tub2-Δ438Q* = 35.3 ± 11.4 nm/s, p = 0.0512; *tub2-Δ438* = 39.5 ± 21.4 nm/s, p = 0.1346 compared to wild type; Supplemental Figure 3). We did not observe any change in Cin8 directional switching (Supplemental Figure 3). Overall, these results indicate that the β-CTT acidic patch promotes midzone-directed Cin8 velocity *in vivo* and plus-end directed velocity *in vitro*.

### The Cin8 N-terminal Extension Interacts with the β-CTT

Because the negative charge of the β-CTT promotes Cin8 localization and motility, we predicted that a positively charged region within Cin8 may mediate this interaction. To objectively identify regions of interest we grouped the kinesin motor domains by secondary structure and sequence alignment from the amino-terminal start through the neck linker at the carboxy-terminal end. We then computationally calculated the isoelectric point for these different regions (Figure 6A). We compared Cin8 and Kip1 to the human kinesin-5 KIF11/EG5 and the ubiquitous kinesin-1 KIF5B. We identified three positively charged regions of interest for Cin8: the amino-terminal extension, loop 8, and the neck-linker. The region containing loop 8 also includes β5 because loop 8 interrupts the two β5 sheets. We chose to focus on the N-terminal extension and loop 8 because both these regions have been previously indicated to play a role in kinesin-5 motility (Britto et al., 2016; Gerson-Gurwitz et al., 2011; Goulet et al., 2014; Pandey, Reithmann, et al., 2021; Singh et al., 2024). Furthermore, both regions have key differences between Cin8 and Kip1, potentially explaining their divergent behaviors: the N-terminus of Cin8 lacks two potential Cdk1 phosphorylation sites found in Kip1, and the loop 8 of Cin8 contains a large 99 residue insertion (Chee & Haase, 2010; Hoyt et al., 1992).

**Figure 6:**
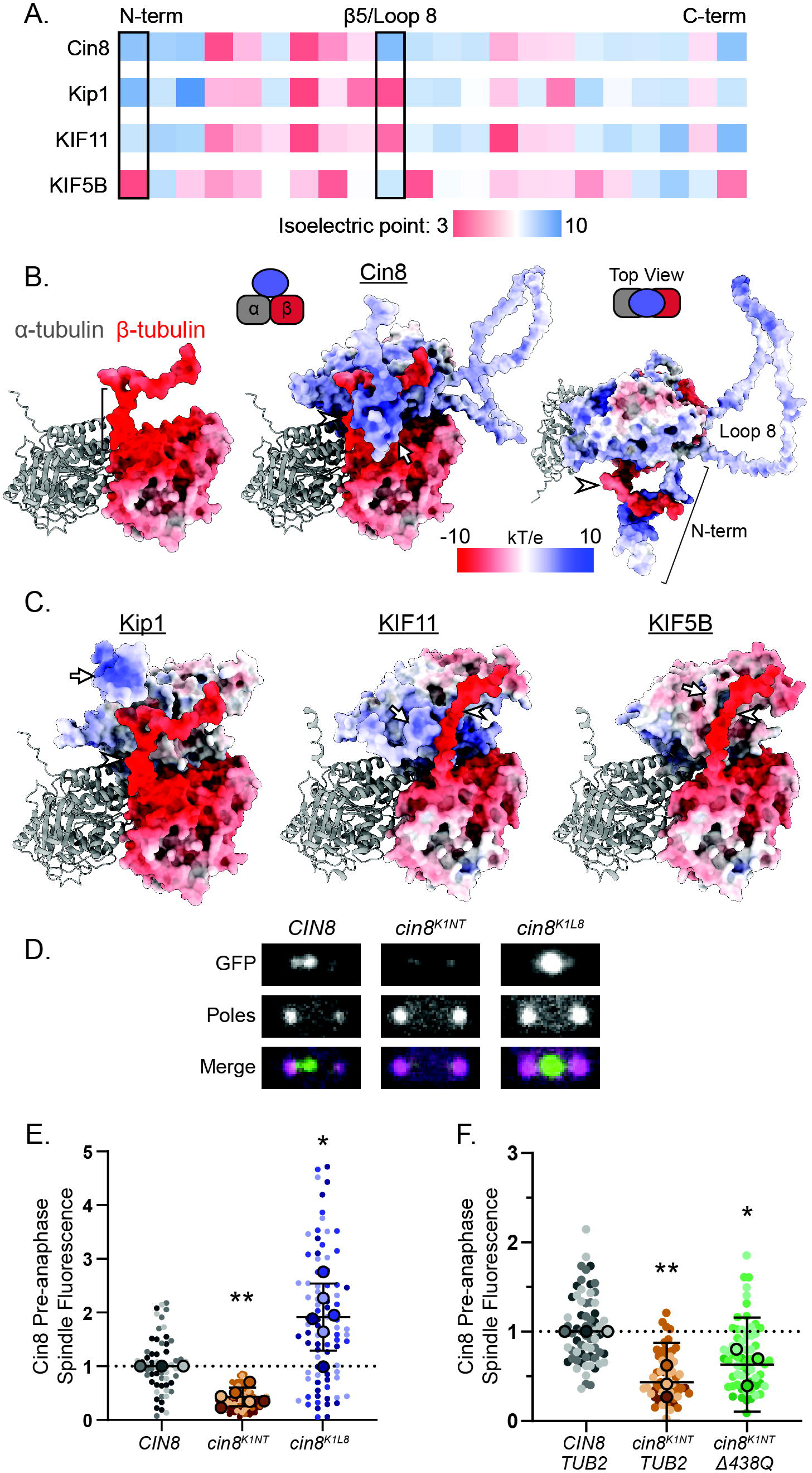
The Cin8 N-terminal Extension Interacts with the β-CTT. **(A)** Computational isoelectric point analysis of kinesin motor domains categorized by secondary structure regions from the amino-terminal through the neck linker for Cin8, Kip1, KIF11 (human kinesin-5), or KIF5B (human kinesin-1). **(B)** Image of a heterodimer (Tub1 and Tub2) demonstrating electrostatic surface potential for β-tubulin, including the β-CTT (left, bracket marks acidic patch), or depicting a feasible interaction with the intrinsically disordered Cin8 N-terminal region from a side view (middle). A top view looking down on the heterodimer is shown on the right. Protein surfaces are colored by electrostatic potential calculated by Poisson-Boltzmann solution. Arrow generally marks Cin8 N-term, and arrowhead marks β-CTT. **(C)** Images demonstrating feasible interactions of the intrinsically disordered kinesin N-terminal regions and the β-CTTs. For kinesin motor domains and β-tubulin, protein surfaces are colored by electrostatic potential calculated by Poisson-Boltzmann solution. Arrows generally mark kinesin N-termini and arrowheads mark β-CTT. For Kip1, Tub1 (α-tubulin) and Tub2 (β-tubulin) are modeled. For KIF11 and KIF5B, TUBA1A (α-tubulin) and TUBB5 (β-tubulin) are modeled. **(D)** Example images of pre-anaphase cells expressing Cin8-3GFP, cin8^K1NT^-3GFP, or cin8^K1L8^-3GFP. Poles are labeled with Spc110-tdTomato. Image width = 2.5 μm. **(E)** Quantification of pre-anaphase spindle fluorescence of cells expressing wild-type Cin8-3GFP (n = 61 cells), cin8^K1NT^-3GFP (n = 88 cells, p < 0.001), or cin8^K1L8^-3GFP (n = 84, p = 0.037). **(F)** Quantification of pre-anaphase spindle fluorescence of cells expressing wild-type Cin8-3GFP (n = 68 cells), cin8^K1NT^-3GFP (n = 62 cells, p = 0.005), or cin8^K1NT^-3GFP and tub2-Δ438Q (n = 65 cells, p = 0.039). For E and F, values are normalized to the median wild-type value. Bolded, outlined points represent the median for each replicate. Statistics are student’s t-test of replicate medians compared to wild-type Cin8. Error bars mean ± 95% CI.

To determine if these regions could possibly interact with the β-CTT, we determined the protein surface electrostatic potential using the Adaptive Poisson-Boltzmann Solver (Figure 6B,C) (Jurrus et al., 2018; Unni et al., 2011). Because these kinesin regions and the β-CTT are intrinsically disordered, we created a potential snapshot of dynamics by using AlphaFold Multimer and then further refinement using “Model loops” in ChimeraX (Cianfrocco et al., 2017; Evans et al., 2022; Meng et al., 2023; Šali & Blundell, 1993). The large loop 8 insertion of Cin8 could feasibly interact with the β-CTT, but what stands out from these models is the close proximity of the negative β-CTT to the kinesin-5 N-termini, particularly for Cin8 (Figure 6B,C, Videos 1-4). The positive charge of the kinesin-5 N-terminus is conserved across species, but is not conserved in other kinesins, for example kinesin-1/KIF5B (Figure 6C; Supplemental Figure 4). These results suggest the β-CTT may interact with the kinesin-5 N-terminal regions through a charge-charge interaction.

Finally, to test if differences in the N-terminal or loop 8 regions explain the selective regulation of Cin8 by the β-CTT, we replaced the Cin8 N-terminus (*cin8 ^K1NT^*) or loop 8 (*cin8 ^K1L8^*) with the corresponding region of Kip1 and quantified the pre-anaphase spindle localization of the fluorescently tagged mutants (Figure 6D,E). Normalized to wild-type *CIN8* cells, cin8 ^K1NT^ exhibited an approximate 50% decrease in spindle localization (mean = 0.43 ± 0.17, p < 0.001), similar to the decrease observed when deleting the β-CTT. Surprisingly, swapping the loop 8 of Kip1 into Cin8 increased spindle localization (cin8^K1L8^ mean = 1.91 ± 0.623, p = 0.037). If the Cin8 N-terminus is interacting with the β-CTT, we predicted that disrupting both the Cin8 N-terminus and the β-CTT would not have an additive effect disrupting Cin8 spindle localization. In agreement with this prediction, double mutant *cin8 ^K1NT^ tub2-Δ438Q* cells (mean = 0.63 ± 0.52, p = 0.039 relative to wildtype; Figure 6F) did not further disrupt localization relative to *cin8 ^K1NT^* alone (p = 0.29), and if anything might slightly increase cin8 ^K1NT^ localization from additional mitotic delay. These results, together with the computational model, suggest that the positive Cin8 N-terminus interacts with the negative β-CTT to promote spindle localization.

## Discussion

In this study, we show that the two budding yeast kinesin-5 motors Cin8 and Kip1 exhibit selective sensitivity to the β-CTT. Our study uses genetic manipulation of the β-CTT to maintain native protein levels and determine the *in vivo* consequences of altering CTT-kinesin interactions and an *in vitro* system to quantify these effects on Cin8 by its native yeast tubulin substrate. Together, our results provide molecular insights into how the β-CTT regulates Cin8, how this affects kinesin-5 function, and suggest a general mechanism for how tubulin code may be functionally read and used to modulate the activity of different kinesins.

A key prediction of the tubulin code hypothesis is that microtubule-associated proteins and motors should exhibit differing sensitivities to the tubulin CTTs. We identified components that were both insensitive (Kar3 and Ase1) and sensitive (Kip3, Cin8, and Kip1) to the loss of the β-CTT (Figure 1). Most strikingly, the two kinesin-5 motors exhibit opposite responses to the loss of the β-CTT, with Cin8 spindle localization increasing and Kip1 decreasing. We attribute this result to the β-CTT directly promoting Cin8 function, which in turn promotes efficient mitotic progression and prevents the accumulation of Kip1 (Figure 3G). Several observations further support our proposed model. First, the Kip1 increase observed in *tub2-Δ430* cells (61%, Figure 1F) is similar to the increase in *cin8Δ* cells (57%, Figure 3C), suggesting loss of Cin8 function is sufficient to increase Kip1 spindle localization. Second, when wild-type cells are arrested for two hours with hydroxyurea, spindles have simultaneously elevated levels of Cin8 and Kip1 (Figure 3E,F), which would not agree with a purely competition-based model. Finally, even when Cin8 spindle localization is decreased to different extents in *tub2-Δ430* and *tub2-Δ438Q* (respectively 41% and 14% decreases; Figure 4B), either disruption of Cin8 localization results in a similar extent of Kip1 enrichment (respectively 56% and 76% increases; Supplemental Figure 2). Altogether these data support a model in which the β-CTT acidic patch directly promotes the localization of Cin8 to the spindle. Kip1 does not appear to be directly regulated by β-CTT during pre-anaphase but is indirectly affected by the loss of Cin8 and consequential delay in mitotic progression. This exemplifies an underappreciated aspect of the tubulin code hypothesis: tubulin CTTs may balance motor and MAP activity through direct and indirect mechanisms. This is an important and cautionary point for future investigations of the tubulin code.

Our results support a model in which electrostatic interactions between the β-CTT acidic patch and the Cin8 basic N-terminus promote Cin8 recruitment to the spindle and plus-end directed motility toward the midzone (Figure 7A,B). Cin8 spindle levels peak prior to anaphase onset, and this localization is supported by the acidic patch of the β-CTT (Figure 2D,4B). Replacing the Cin8 N-terminus with that of Kip1 results in an ∼50% reduction in spindle localization (Figure 6E), phenocopying the genetic deletion of the β-CTT. This interaction between the β-CTT and Cin8 N-terminus is supported by their opposite charges and close proximity (Figure 6B,C), and recent structural work from other labs (Singh et al., 2024). Our computational isoelectric point calculations might suggest that the Cin8 and Kip1 N-termini should behave similarly (Figure 6A, Supplemental Figure 4). However, it is important to note that phosphorylation of unique sites in the Kip1 N-terminus may explain the divergent behavior, and provide a possible point of Kip1 regulation (Chee & Haase, 2010). Resolving these possibilities and the regulation of the two budding yeast kinesin-5 motors will require further study.

**Figure 7:**
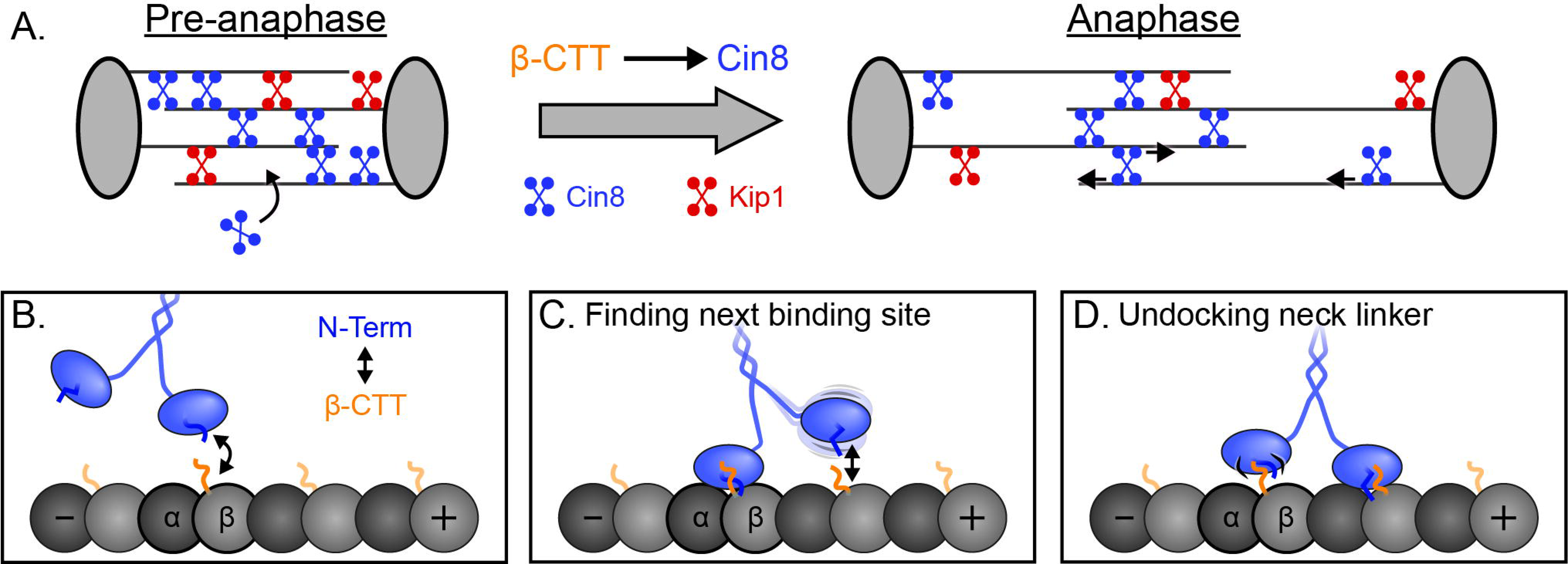
Model – The β-CTT interacts with the kinesin-5 N-terminus to promote motility. **(A)** Spindle localization of kinesin-5 reaches maximum prior to anaphase onset with higher levels of Cin8 relative to Kip1. The β-CTT increases Cin8 spindle localization. During spindle elongation in anaphase, the β-CTT increases Cin8 motility and promotes midzone localization. **(B)** Interactions between the β-CTT and Cin8 N-terminus increase Cin8 localization to microtubules. **(C)** Then, during the Cin8 mechanochemical cycle, this interaction could promote the binding of the forward motor domain, increasing the Cin8 velocity. **(D)** Finally, this interaction could help release the trailing motor domain as well as maintain association between the motor domain and microtubules within the midzone during spindle elongation.

The interaction between the β-CTT and Cin8 N-terminus is also important for specifically localizing Cin8 to the spindle midzone, which we attribute to increased plus-end directed motility. We find that the acidic patch of the β-CTT is sufficient to increase Cin8 plus-end velocity, both *in vivo* and *in vitro* (Figure 5). Using an *in vitro* system we developed to study Cin8 motility on genetically modified yeast tubulin, we quantified an increase in Cin8 plus-end directed run length with the β-CTT acidic patch, but no change in run duration. How might these effects be related to the mechanochemical cycle of Cin8? We propose two steps in the mechanochemical cycle that could be promoted by interaction between the Cin8 N-terminus and β-CTT: 1) the leading motor domain finding the next binding site along the microtubule (Figure 7C) or 2) release of the trailing motor domain to enable the next forward step (Figure 7D). Past biochemical work with *X. laevis* KIF11 suggested that finding the next binding site is one of the fastest steps in the kinesin-5 mechanochemical cycle, while release of the trailing motor domain is the rate limiting step (Chen et al., 2016). These experiments with KIF11 were conducted with brain-derived tubulin, which is enriched for glutamylation of the tubulin CTTs; therefore, the fast rate of binding to the next site may be in part because of enhanced electrostatic interactions between the KIF11 N-terminus and β-CTT. Regarding release of the trailing head, cryo-electron microscopy structures of human KIF11 identified a partial density for the N-terminus that underwent conformational changes coupled with the mechanochemical cycle of kinesin-5 (Goulet et al., 2014). Release of the N-terminal cover strand interaction with the neck-linker allowed for undocking of the neck-linker from the kinesin motor domain to reset for the next step (Goulet et al., 2014). A similar density was recently observed for a structure of AMPPNP-bound Cin8, and the N-terminus of Cin8 was necessary for Cin8 plus-end motility (Singh et al., 2024). Interactions between the β-CTT and the kinesin-5 N-terminus could support these transitions during the mechanochemical cycle to promote kinesin-5 motility.

It is important to note that our *in vitro* experiments measure Cin8 motility on single microtubules, and do not examine the antiparallel microtubule context that is key to kinesin-5 tetramer function in the mitotic spindle. Research with both human KIF11 and *D. melanogaster* KLP61F demonstrated that in the tetramer, kinesin-5 carboxy-tail domains of one pair of motor domains interact with the other pair of motor domains to alter the mechanochemical cycle and sustain the sliding of antiparallel microtubules (Bodrug et al., 2020). These interactions occur through an α0-helix connected to the N-terminal region and could thus influence the kinesin-5 N-terminus (Bodrug et al., 2020). Our *in vivo* results suggest the β-CTT is important for maintaining Cin8 at the spindle midzone (Figure 4). As Cin8 slides apart antiparallel microtubules while experiencing resistive loads, the interaction between Cin8 and the β-CTT to promote motility may be even more important to contribute to spindle elongation.

We propose that interaction between the N-terminus and the β-CTT is a conserved mechanism of kinesin-5 motors that may enable selective regulation of motor activity within the spindle. This model is supported by our electrostatic surface analysis and the broad conservation of the positive charge of kinesin-5 N-termini (Figure 6, Supplemental Figure 4). Moreover, the N-terminal regions of *S. pombe* Cut7 and human KIF11 have both been demonstrated to be important for their motility (Britto et al., 2016; Goulet et al., 2014). An attractive feature of our model is that regulation could be bilateral in some organisms, acting through kinesin or β-CTT or both. For budding yeast, there is only a single β-tubulin isotype and no known tubulin PTMs, so regulation is unilateral and occurs via the different kinesin-5 motors or via kinesin phosphorylation, such as in the N-terminus of Kip1 (Chee & Haase, 2010). In metazoans, polyglutamylation of the β-CTT further increases its negative charge and potential interactions with kinesin N-terminal regions. Polyglutamylation is enriched on mitotic spindle microtubules where kinesin-5 functions (Bobinnec et al., 1998). How modifications of kinesin-5 N-termini and tubulin CTTs converge to control mitotic spindle function, and whether similar regulation extends to other kinesins will be important topics for future studies.

## Methods

### Yeast strains and manipulation

Chemicals and reagents purchased from Thermo Fisher Scientific (Waltham, MA) and Sigma-Aldrich (Saint Louis, MO), unless stated otherwise. General yeast manipulation, media, and transformation were standard protocols. Fluorescent tag fusion to Ase1, Kar3, Kip1, Kip3, and Spc110 are integrated at the corresponding native loci. 3HA affinity tag fusion to Cin8 and Kip1 are at their corresponding native loci. Bik1-3GFP integrating plasmid was a gift from Dr. David Pellman (Harvard University) and integrated at the endogenous locus. The Cin8-3GFP integrating plasmid targets to the endogenous *CIN8* locus, and the same integrated allele was used for all experiments. The mNeonGreen plasmid DNA was provided by Allele Biotechnology and Pharmaceuticals (San Diego, CA) (Shaner et al., 2013). Mutant alleles of *tub2* and *cin8* were generated at the endogenous loci. All mutations were confirmed by Sanger sequencing of the genomic loci, and were confirmed as the only mutations present in the coding sequence. Yeast strains are listed in Supplementary Table 1.

### Live cell imaging

Cells were grown overnight in rich media (2% glucose, 2% peptone, and 1% yeast extract) at 30°C, diluted into fresh rich media and grown to log phase at 30°C, then washed and imaged in synthetic media (2% glucose, CSM from Sunrise Science Products, #1001 San Francisco, CA). Cells were adhered to slide chambers coated with concanavalin A and sealed with VALAP (Vaseline, lanolin, and paraffin at 1:1:1) (Fees et al., 2017). Images were collected on a Nikon Ti-E microscope equipped with a 1.45 NA 100× CFI Plan Apo objective, piezo electric stage (Physik Instrumente, Auburn, MA), spinning disk confocal scanner unit (CSU10; Yokogawa), 488-nm and 561-nm lasers (Agilent Technologies, Santa Clara, CA), and an EMCCD camera (iXon Ultra 897; Andor Technology, Belfast, UK) using NIS Elements software (Nikon). All live cell imaging conditions were as follows unless otherwise noted: samples were maintained at 30°C using an ASI 400 Air Stream Incubator (NEVTEK, Williamsville, VA). Each image was a z-stack consisting of 13 images separated by 450 nm.

### Spindle localization analysis

Pre-anaphase cells were identified and imaged from an asynchronous log phase population by only observing Spc110-tdTomato signal. The z-stack consisting of 13 images separated by 450 nm was then sum projected for analysis and pre-anaphase cells were considered to have a spindle length < 2 µm. The spindle ends were defined as the Spc110-tdTomato signal, and a line with a width of 5 pixels (∼300 nm) was drawn around this region to determine the fluorescence intensity on the spindle. Background subtraction was performed by drawing a 13-by-13 pixel box (0.481 µm^2^) in the cytoplasm to determine the average cellular background fluorescence and then normalized to the spindle area and subtracted from the spindle intensity. Values were normalized to the median value of the two wild-type biological replicates for each of the three technical replicates.

### Kinesin-5 spindle localization timecourse

Log phase cells were arrested with alpha factor for 3 hours and then released and grown at 30°C and imaged starting 60 minutes after release. Each z-stack image was taken every 2 minutes. Z-stacks were sum projected, and the spindle endpoints were defined by Spc110-tdTomato signal with a 5-pixel wide segmented line drawn between these two endpoints along the spindle estimated by kinesin-5 fluorescence. Background subtraction was performed by drawing a 13-by-13 pixel box in the cytoplasm to determine the average cellular background fluorescence and then normalized to the spindle area and subtracted from the spindle intensity. Values were normalized to the start of anaphase onset as determined by the timepoint immediately preceding a rapid change in spindle length.

### Anaphase spindle localization and recruitment

Anaphase cells were identified and imaged from an asynchronous log phase population by only observing Spc110-tdTomato signal. Anaphase cells for analysis were considered to have a spindle length between 2.5 and 6 µm. The imaged z-stack was sum projected for analysis, and the spindle endpoints were defined by Spc110-tdTomato signal with a 3-pixel wide segmented line drawn between these two endpoints along the spindle estimated by Cin8-3GFP fluorescence. The average intensity of the 3 pixels at each point of the line was used to determine the fluorescence intensity at that point along the spindle. Background subtraction was performed by drawing a 13-by-13 pixel box in the cytoplasm to determine the average cellular background fluorescence per pixel and this average value was subtracted from each point along the spindle. The total anaphase Cin8-3GFP fluorescence was the total background-subtracted intensity values along the spindle line. The spindle length was then divided into five equal parts and the sum fluorescence intensity along that fifth of the spindle length was divided by the total anaphase fluorescence to determine the proportion of signal in that region of the spindle. Total anaphase intensity values were normalized to the median value of the two wild-type biological replicates for each of the three technical replicates.

### *In vivo* Cin8 motility

Early to mid-anaphase cells were identified and imaged from an asynchronous log phase population by observing Spc110-tdTomato signal. Each z-stack was then taken every 5 seconds. Kymographs were generated by drawing a 10-pixel wide segmented line along the spindle, and then the “Straighten” function was used to generate a centered image stack from which the kymographs were made. Motility events were objectively identified using KymoButler with default settings (threshold 0.20, minimum size: 3, minimum frames: 3) (Jakobs et al., 2019), and the first and last points were used to calculate the average velocity.

### *In vitro* Cin8 motility on yeast microtubules

One-liter cultures of yeast were grown to log phase in rich media, pelleted by centrifuging at 4,000 g at 4°C, and then washed twice with water. Enough 10X BRB80 (80 mM PIPES, 1 mM MgCl_2_, 1 mM EGTA) with 150 mM KCl was added to pipette the pellet (approximately 500 µl per 5 ml of yeast pellet), and the yeast slurry was snap frozen by pipetting single drops into liquid nitrogen and stored at −80°C. The cells were lysed by grinding in 50 ml grinding jar chilled in liquid nitrogen in a Mixer Mill MM 400 (Retsch, Hamburg, Germany). Two cycles of 30 Hz for 110 seconds were used, allowing the jars to chill in liquid nitrogen after each run. Powdered lysate was then collected in a chilled 50 ml conical and stored at −80°C.

Reaction chambers were assembled with a plasma cleaned and HMDS silanized (Wedler et al., 2022) 22-by-22 mm coverslip and an 18-by-18 mm coverslip between which were melted single ply strips of parafilm to create two chambers on a custom-made stage insert. Anti-rhodamine antibody (Thermo Fisher A-6397) was diluted 1:100 in cold BRB80 and flowed into a chamber to incubate for 5 min at room temperature, and then the chamber was then washed with 100 µl of BRB80. 50 µl of 1% pluornic-F127 in BRB80 was flowed into the chamber to incubate for 5 min at room temperature, and then washed with 100 µl of BRB80. Rhodamine-labeled GMPCPP-stabilized porcine brain microtubule seeds were then flowed into the chamber for 30 s before washing out with 200 µl of BRB80 and were then ready for use.

This protocol was based on a protocol from the Barnes lab (Bergman et al., 2018; Torvi et al., 2022). The yeast lysate powder was prepared as follows: two polypropylene ultracentrifuge tubes were chilled on ice and a small amount (< 100 mg) of prepared yeast lysate powder was added to each tube directly from the −80°C freezer. Each individual tube was weighed before and after addition of the powdered lysate to determine the specific amount. One tube was used to make a “high concentration” clarified lysate using a ratio of 0.1 µl of 10X BRB80 plus 150 mM KCl per mg of powdered lysate; the other tube was used to make a “low concentration” of clarified lysate using a ratio of 2 µl of 10X BRB80 plus 150 mM KCl per mg of powdered lysate. Additionally, 1 µl of yeast protease inhibitor cocktail (Sigma-Aldrich P8215) was added to each preparation. The tubes were then briefly vortexed and spun down in a desktop centrifuge to gather everything at the bottom of each tube and the lysate was fully thawed on ice for 10 minutes. The powdered lysate was then clarified by centrifugation at 100,000 g for 30 minutes at 4°C, and the supernatant was transferred to a chilled microcentrifuge tube for use in the reactions. The high concentration clarified lysate was approximately 7.5 mg/ml protein and the low concentration was approximately 4 mg/ml, as measured by Pierce 660 nm protein assay reagent.

To first polymerize microtubules, a 50 µl reaction consisting of 10 µl of the high concentration clarified yeast lysate (final protein concentration of approximately 1.5 mg/ml) in BRB80 containing 1 mM MgCl_2_, 8 mM ADP, 1 mM GTP, and 2 µM epothilone A was flowed into a prepared chamber. Yeast microtubules polymerized for 10 min at 30°C. To remove immotile Cin8-3GFP clusters and other proteins from the microtubules, the chamber was then washed twice with 50 µl of 750 mM KCl, 100 µM epothilone A in BRB80 for 3 minutes each, and then once for 3 minutes with 100 µl of BRB80, all maintaining 30°C. A 50 µl reaction was then prepared with 0.5 µl of the low concentration clarified lysate (final protein concentration of approximately 40 ug/ml) in BRB80 containing 1 mM MgCl_2_, 2 mM ATP, 1 mM GTP, and 100 µM of epothilone A and flowed into the chamber and sealed with VALAP for imaging at 30°C.

Images were collected on a Nikon Ti-E microscope equipped with a 1.49 NA 100× CFI160 Apochromat objective, TIRF illuminator, OBIS 488-nm and Sapphire 561-nm lasers (Coherent, Santa Clara, CA), W-View GEMINI image splitting optics (Hamamatsu Photonics, Hamamatsu, Japan) and an ORCA-Flash 4.0 LT sCMOS camera (Hammamatsu Photonics). Stage was heated to 30°C using an ASI 400 Air Stream Incubator (NEVTEK, Williamsville, VA). Microtubules were verified by interference reflection microscopy. Images were collected by TIRF imaging at 2 second intervals. Cin8 remained motile through 90 total minutes of imaging. Kymographs were generated by drawing a 10-pixel wide segmented line along a microtubule, and then the “Straighten” function was used to generate a left-aligned image stack from which the kymographs were made. Kymographs were analyzed by identifying the first and last points of the motility event and calculating the displacement and duration between these two points. KymoButler with default settings (threshold 0.20, minimum size: 3, minimum frames: 3) was used to objectively call unclear events (Jakobs et al., 2019).

### Western blotting timecourse

Cells were arrested with alpha factor for 3 hours and then released and samples collected every 15 minutes. Cells were pelleted and resuspended in 2 M lithium acetate and incubated for 3 min at room temperature. Cells were pelleted again and resuspended in cold 0.4 M NaOH for 3 min on ice. Cells were pelleted and resuspended in 2.5X Laemmli buffer and boiled for 5 minutes. Samples were run on 4-15% Mini-PROTEAN TGX precast protein gels (Bio-Rad Laboratories, Hercules, California) in running buffer (250 mM Tris, 1.9 M glycine, 1% SDS at pH 8.3) for 100 V for 10 min and then 130 V for one hour. Gels were transferred to PVDF (Millipore, IPFL85R) in transfer buffer (250 mM Tris, 1.9 M glycine, 10% methanol) at 20V for 13 hours at 4°C. Membranes were blocked for 1 hour at room temperature in PBS blocking buffer (LI-COR, 927-70001). Membranes were probed in PBS blocking buffer with mouse-anti-HA antibody (Santa Cruz SC-7392) at 1:500 and rabbit-anti-Zwf1 (Sigma A9521) at 1:10,000 for 3 hours at room temperature and then washed three times in PBS at room temperature. Membranes were probed with secondary antibodies goat-anti-mouse-680 (LI-COR 926-68070) and goat-anti-raabbit-800 (LI-COR 9226-32211) both at 1:15,000 in PBS blocking buffer for 1 hour at room temperature. After incubation, membranes were washed three times in PBS with 0.1% Tween-20, three times in PBS, and then imaged on an Odyssey Imager (LI-COR 2471). Images were analyzed in FIJI using the Analyze > Gels function.

### Doubling time assays

Cells were grown to saturation in rich liquid media at 30°C and then diluted 50X into fresh rich media. The diluted cultures were then aliquoted into a 96-well plate and incubated at 30°C with single orbital shaking in a Cytation3 plate reader (BioTek, Winooski, VT). The OD600 was measured every 5 minutes for 24 hours and doubling time calculated by fitting an exponential growth curve to the data.

### Isoelectric point and surface electrostatic potential analysis

Kinesin motor domain alignment was run on Clustal Omega multiple sequence alignment (Madeira et al., 2022) and then secondary structure was determined by comparison between EG5 PDB: 6TA4 and KIF5B PDB: 6OJQ. Secondary structures were grouped for cases in which they could not be clearly distinguished, such as the case of loop 8 interrupting the two sheets of β5. The isoelectric point was then computationally calculated with Isoelectric Point Calculator 2.0 (Kozlowski, 2021). To determine surface electrostatic potentials, we first created representative snapshot models of the intrinsically disordered kinesin and tubulin regions using AlphaFold Multimer and further refinement using the “Model loops” MODELLER function in Chimera X (Cianfrocco et al., 2017; Evans et al., 2022; Meng et al., 2023; Šali & Blundell, 1993). Models were picked that had minimal normalized Discrete Optimized Protein Energy scores and reduced steric clashes. For Cin8 and Kip1 models, Tub1 and Tub2 were used for α- and β-tubulin, respectively. For KIF11 and KIF5B models, TUBA1A and TUBB5 were used for α- and β-tubulin, respectively. Surface electrostatic potential was then calculated by using the PDB2PQR webserver to prepare the PDB file (default settings: PROPKA to assign protonation states at pH 7.0 and PARSE forcefield) and then the Adaptive Poisson-Boltzmann Solver webserver (default settings: “mg-auto”) to calculate the electrostatic surface continuum (Jurrus et al., 2018; Unni et al., 2011). The surface color for electrostatic potential was then applied in ChimeraX (Meng et al., 2023).

### Statistics

Prism (GraphPad Software; San Diego, CA) was used for all graphs and statistical analysis. For all multiple comparisons, an ANOVA test was first performed, and subsequent analysis was performed if p < 0.05. Student’s t-test was used for all homoscedastic, parametric data. Student’s t-test with Welch’s correction was used for heteroscedastic, parametric data. Mann-Whitney *U* test was used for nonparametric data.

## Supplemental Materials

**Supplemental Figure 1: Quantification of β-CTT effect on Bik1 and whole cell example images**

**(A)** Quantification of Bik1-3GFP spindle localization in the presence of absence of the β-CTT. *TUB2* n = 87 cells, *tub2-Δ430* n = 93 cells, p = 0.23. Values are normalized to the median wild-type value. Bolded, outline points represent the median for each replicate. Error bars are the mean ± 95% CI for the replicate medians. Statistics are student’s t-test for the replicate medians.

**(B)** Example images from Figure 1 showing the entire cell, including Bik1 quantified in (A). The spindle poles are labeled with Spc110-tdTomato. Scale bars = 1 µm.

**Supplemental Figure 2: Pre-anaphase Kip1 spindle localization and full Cin8 anaphase quantification**

**(A)** Quantification of background-subtracted Kip1-mNeonGreen pre-anaphase spindle fluorescence in wild-type *TUB2* or *tub2* mutant cells. Values are normalized to the median wild-type value. Bolded, outlined points represent the median for each replicate. Error bars are mean ± 95% CI for the replicate medians. *TUB2*: n = 105 cells; *tub2-Δ430*: n = 111 cells, p = 0.01; *tub2-Δ438*: n = 120 cells, p = 0.87; *tub2-Δ438Q*: n = 120 cells, p = 0.01. Statistics are student’s t-test with Welch’s correction compared to wild-type *TUB2*.

**(B-E)** Quantification of proportion of Cin8-3GFP fluorescence along anaphase spindles in cells expressing **(B)** *TUB2*, **(C)** *tub2-Δ430*, **(D)** *tub2-Δ438Q*, or **(E)** *tub2-Δ438Q* from Figure 3E-H. Each quintile represents 20% of the total spindle length. Error bars are median ± 95% CI for all cells.

**Supplemental Figure 3: *In vitro* microtubules from whole cell lysate and Cin8 minus-end velocity**

**(A)** Example images of microtubules assembled *in vitro* from whole cell lysate from cells expressing GFP-Tub1 before and after high salt washes.

**(B)** Example images of *in vitro* motility experiments depicting Cin8-3GFP, rhodamine-labeled porcine brain-tubulin GMPCPP stabilized seeds, and yeast microtubules (denoted by arrowheads) visualized by interference reflection microscopy.

**(C)** Quantification of Cin8 minus-end velocity. *TUB2*: n = 75 events; *tub2-Δ438Q*: n = 115 events, p = 0.051; *tub2-Δ438*: n = 116 events, p = 0.13. Statistics are student’s t-test of replicate medians compared to wild-type *TUB2*. Error bars mean ± 95% CI.

**(D)** Percentage of plus-end directed motility events out of all events (plus-end directed, minus-end directed, or diffusive). Each point is a technical replicate.

**(E)** Percentage of plus-end processive motility time out of all time spent in processive motility (plus-end or minus-end directed motility). Each point is a technical replicate.

**Supplemental Figure 4: Conserved positive charge of kinesin-5 N-terminus**

**(A)** Sequences of N-terminus of multiple kinesin-5 motors through the start of the catalytic motor domain core. KIF5B (kinesin-1) is included for comparison. Positive residues are blue. Bolded black residues indicate Cdk1 phosphorylation sites. Sc: *Saccharomyces cerevisiae*; Sp: *Schizosaccharomyces pombe*; Ce: *Caenorhabditis elegans*; Dm: *Drosophila melanogaster*; Xl: *Xenopus laevis*; Mm: *Mus musculus*; Hs: *Homo sapiens*.

**(B)** Computational isoelectric point of N-terminus for kinesin motors in A. Expressed as most predictive model (average of all models).

**Table S1.**
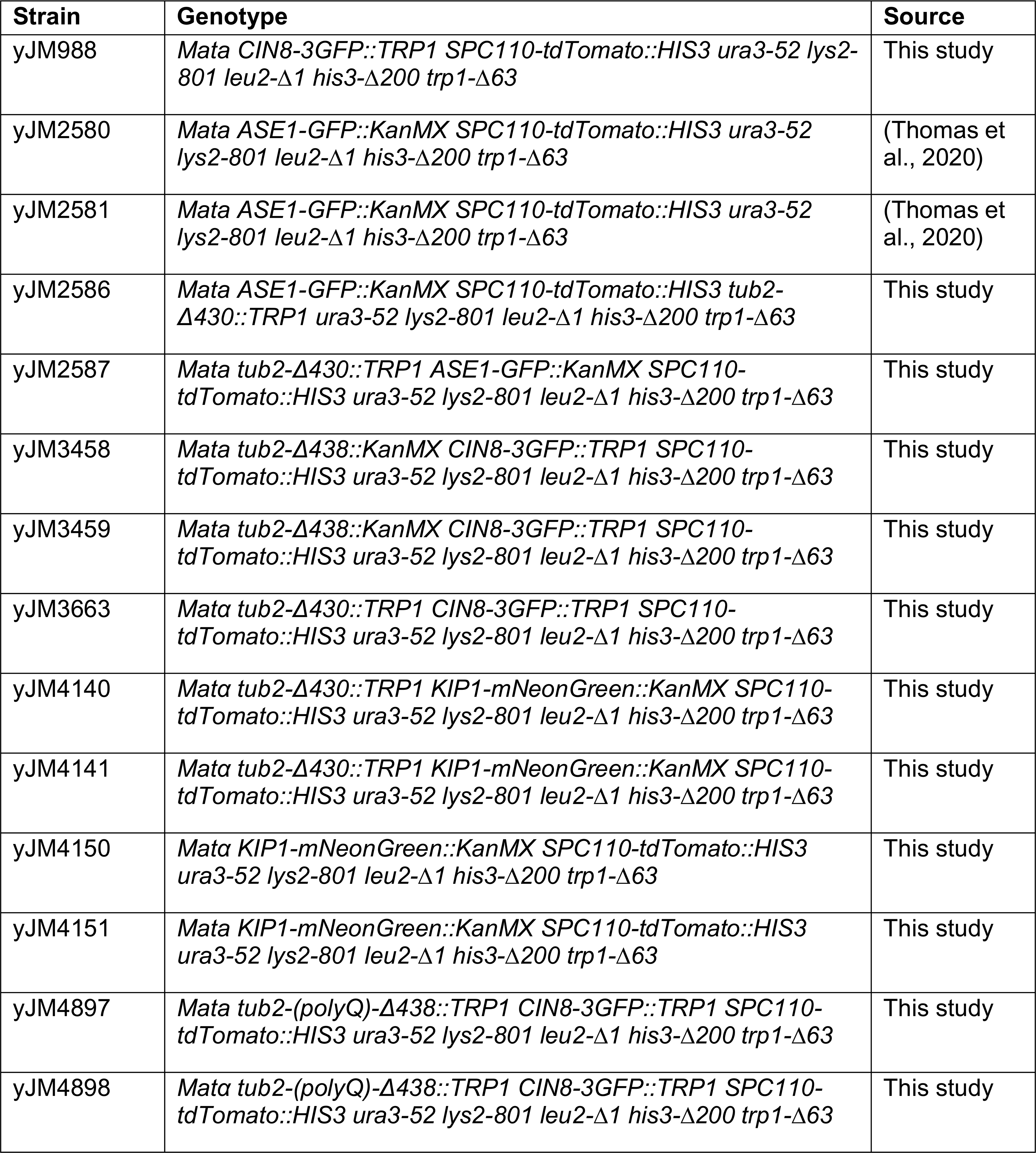
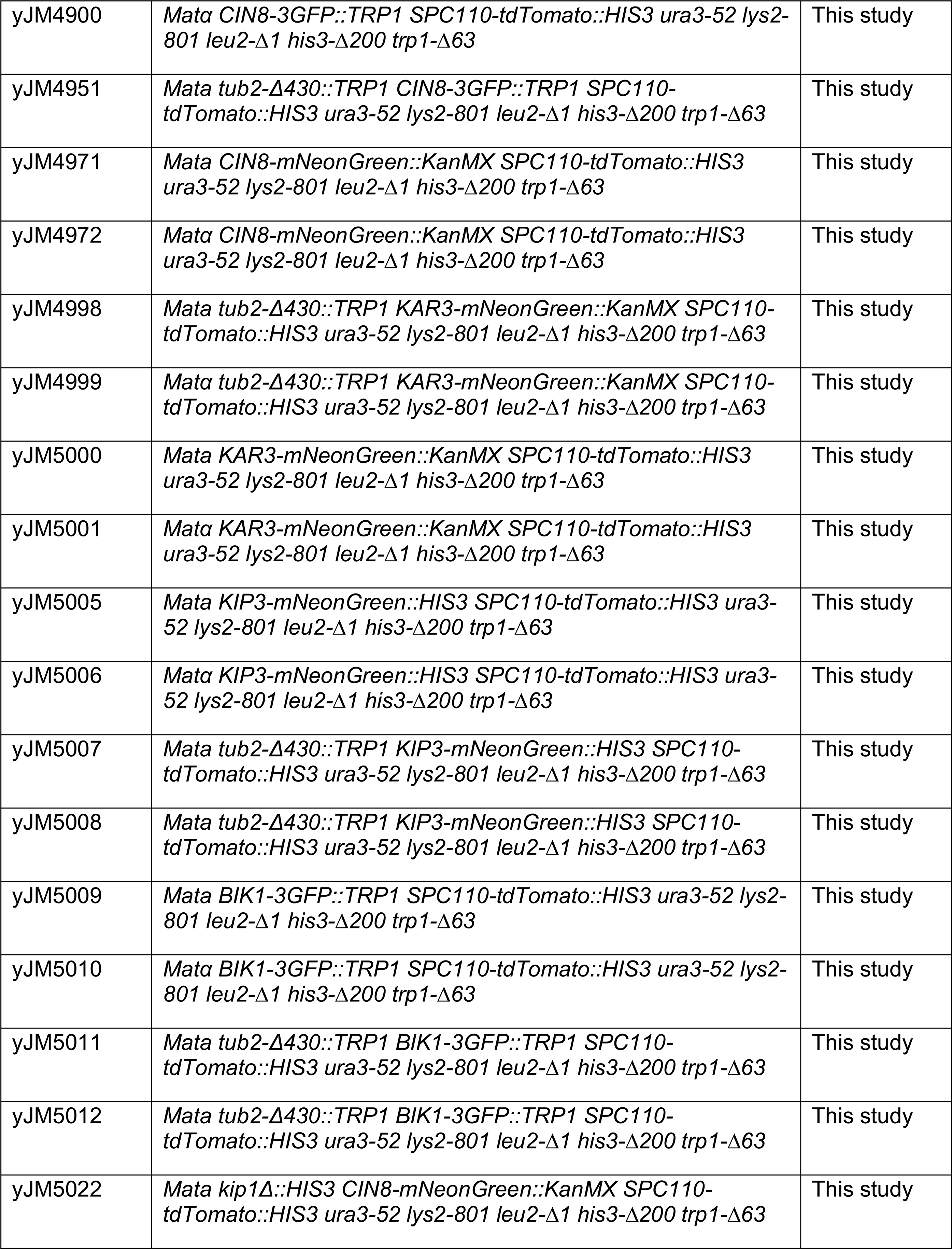
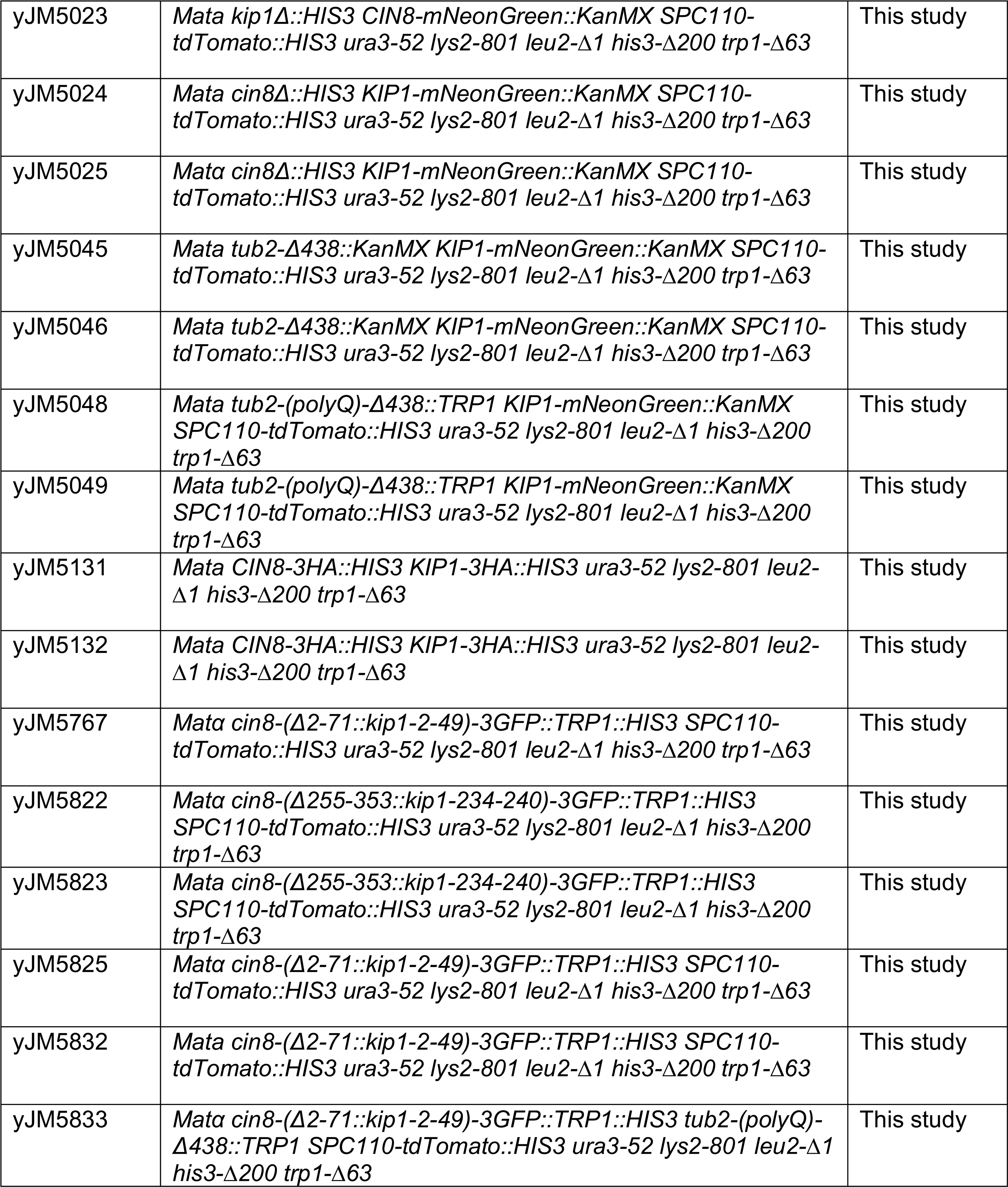
Strains used in this study.

**Videos 1-4:** Depictions of electrostatic surface potential of kinesin motor domain bound to tubulin heterodimer as determined by AlphaFold Multimer with disordered regions further refined by MODELLER (see Methods). Videos 1 and 2 depict Cin8 and Kip1 motor domains, respectively, on Tub1 (α-tubulin) and Tub2 (β-tubulin). Videos 3 and 4 depict KIF11 and KIF5B motor domains, respectively, on TUBA1A (α-tubulin) and TUBB5 (β-tubulin). Alpha tubulin is in grey for every video. Electrostatic surface potential (kT/e) calculated by Poisson-Boltzmann solution.

**Video 1: Electrostatic surface potential of Cin8 motor domain and β-tubulin**

**Video 2: Electrostatic surface potential of Kip1 motor domain and β-tubulin**

**Video 3: Electrostatic surface potential of KIF11 motor domain and β-tubulin**

**Video 4: Electrostatic surface potential of KIF5B motor domain and β-tubulin**

