## Supplementary figures and images for "Selective regulation of kinesin-5 function by β-tubulin carboxy-terminal tails"

### Supplemental Figure 1

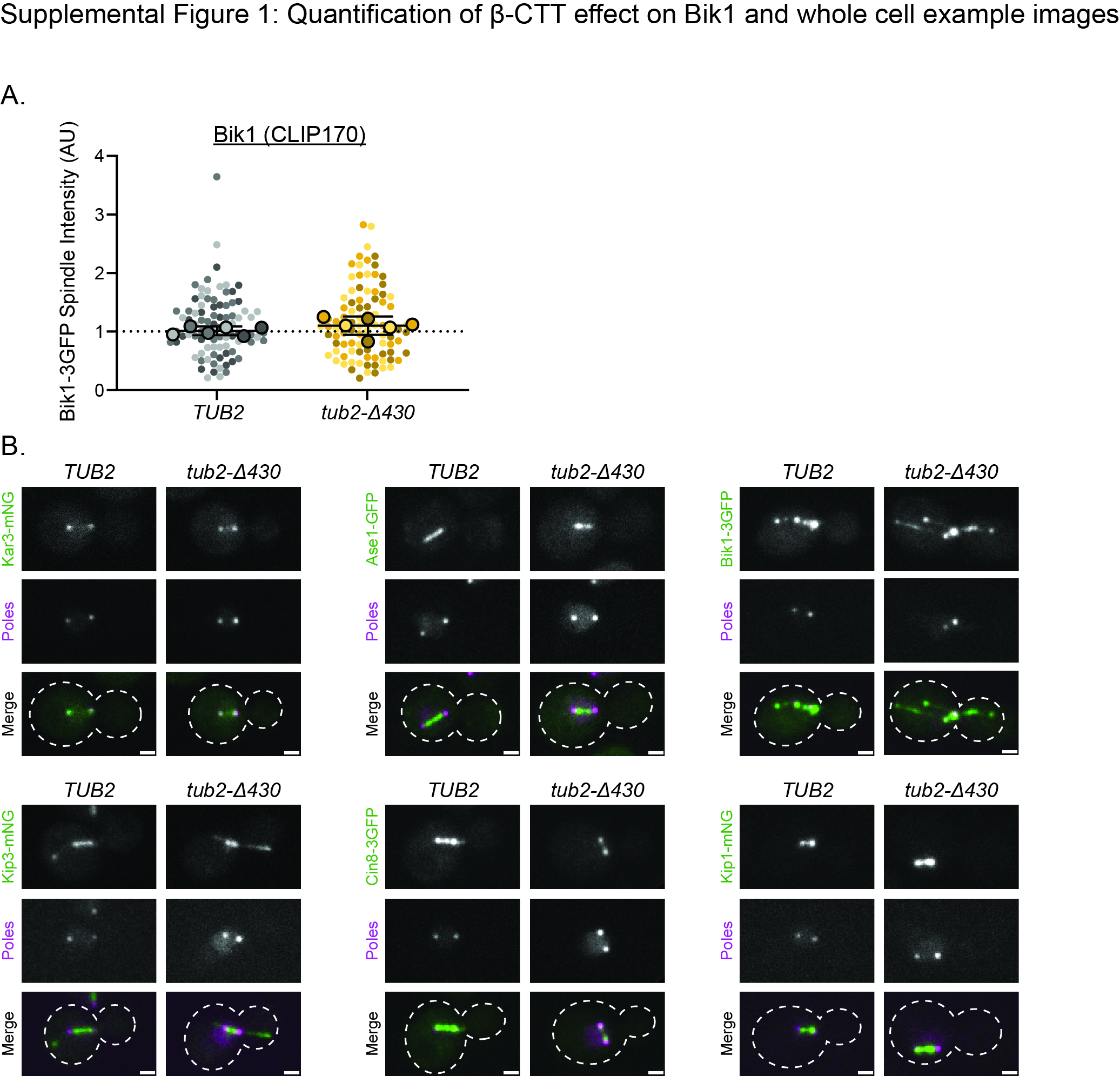

### Supplemental Figure 2

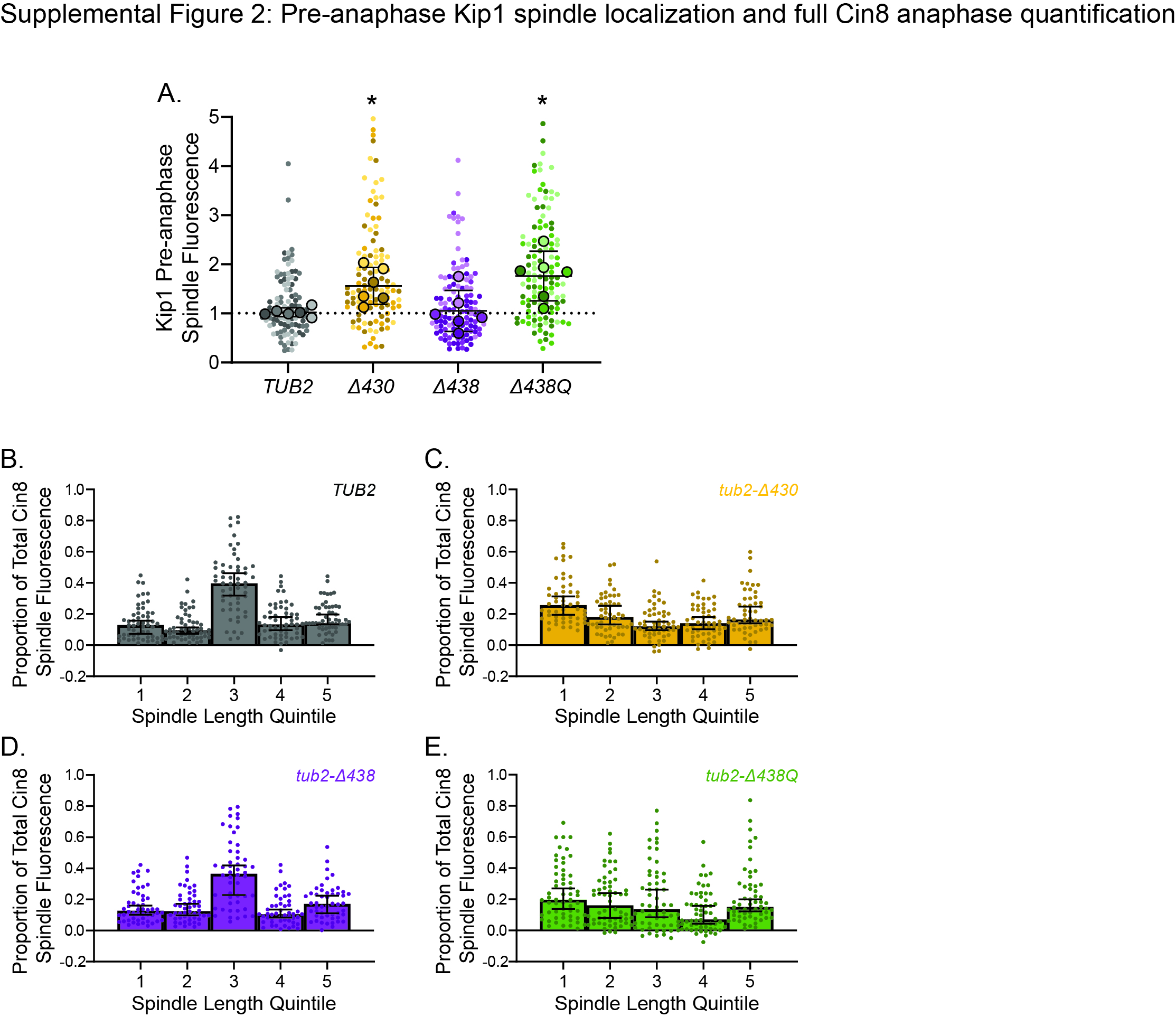

### Supplemental Figure 3

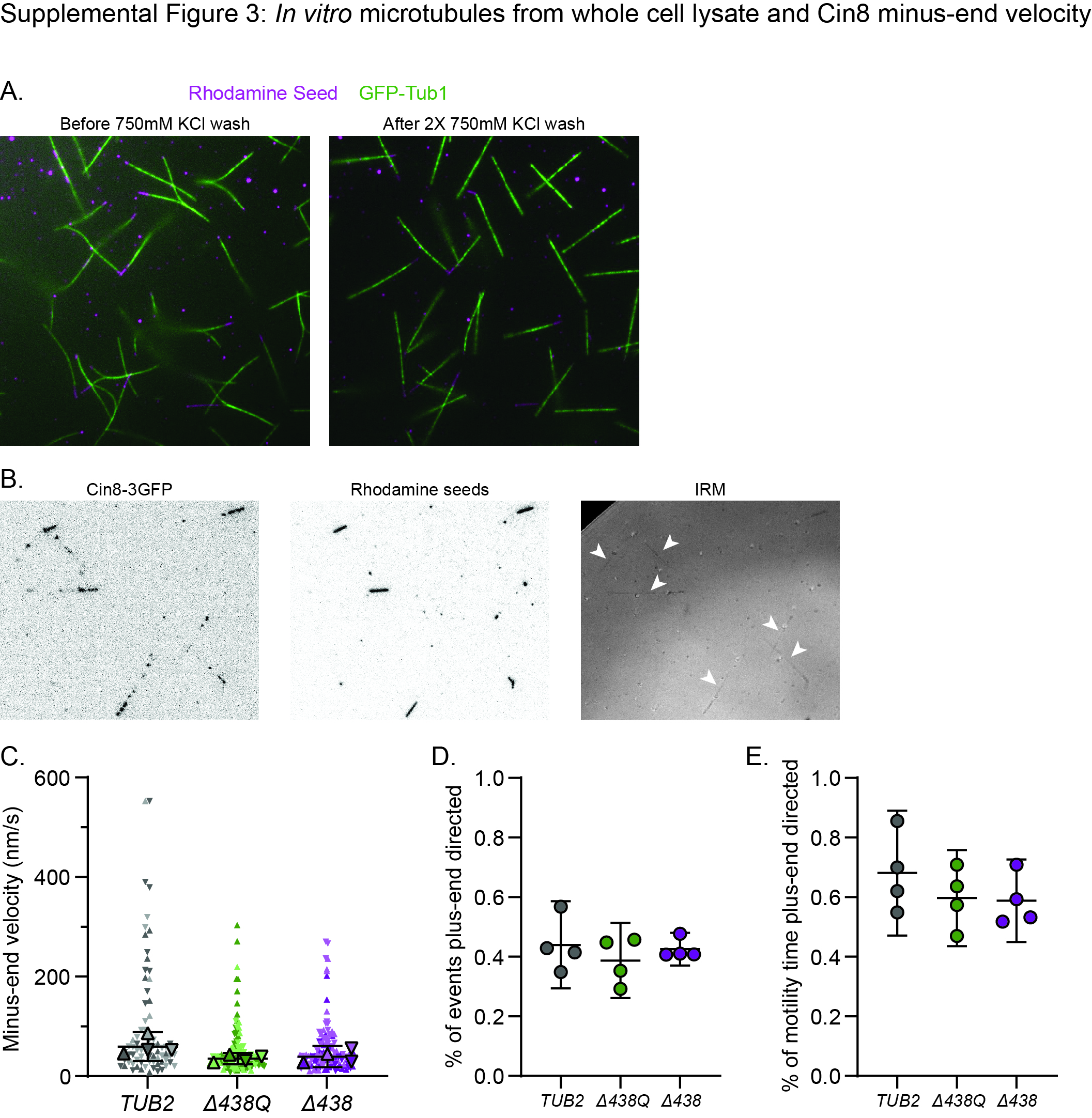

### Supplemental Figure 4

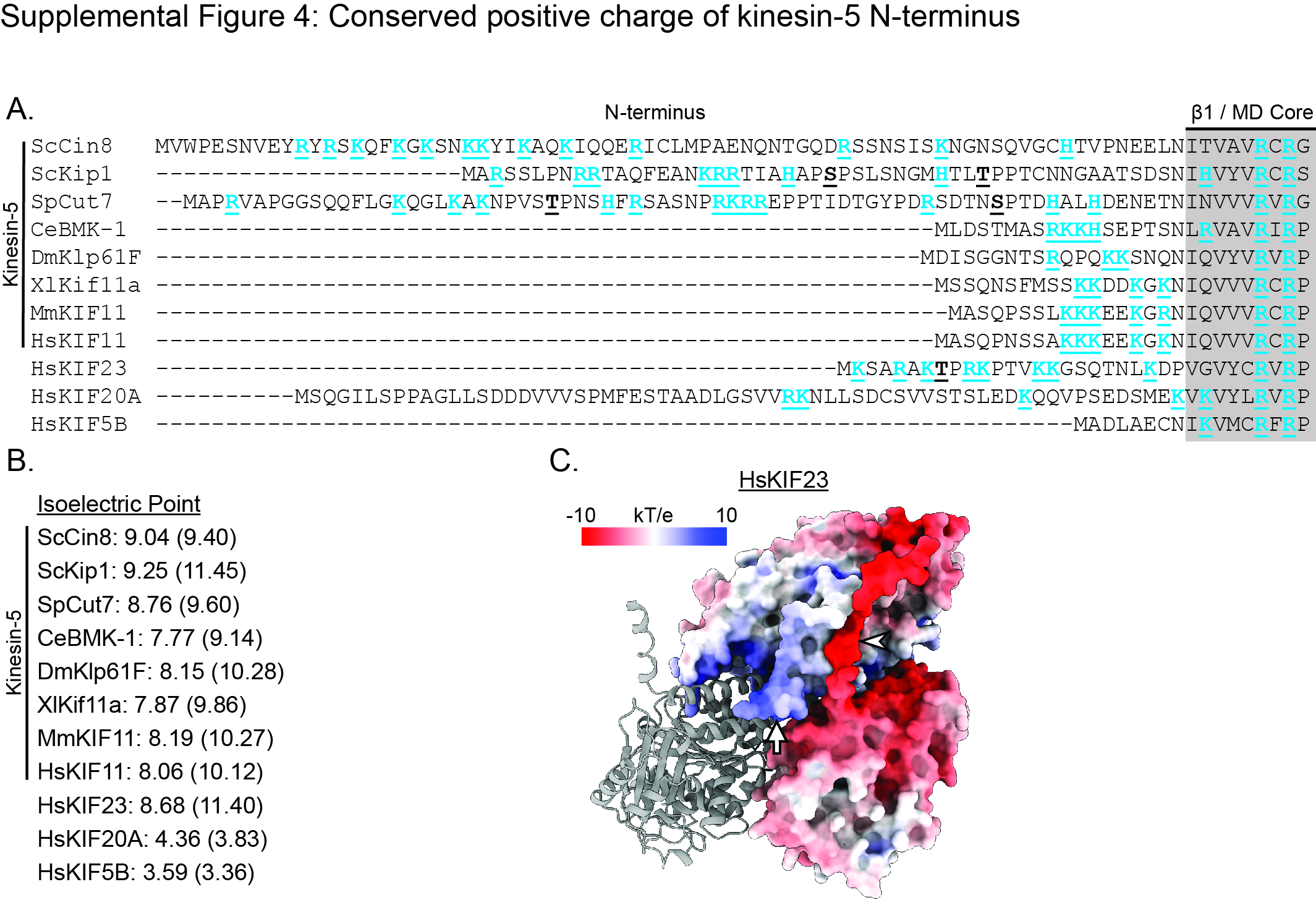
